# Life stage impact on the human skin ecosystem: lipids and the microbial community

**DOI:** 10.1101/2024.01.03.573871

**Authors:** Martin P. Pagac, Bala Davient, Hilbert Yuen In Lam, Aarthi Ravikrishnan, Wee Ling Esther Chua, Sneha Muralidharan, Aishwarya Sridharan, Antony S. Irudayaswamy, Ramasamy Srinivas, Stephen Wearne, Ahmad Nazri Mohamed Naim, Eliza Ho Xin Pei, H. Q. Amanda Ng, Junmei Samantha Kwah, Eileen Png, Anne K. Bendt, Markus R. Wenk, Federico Torta, Niranjan Nagarajan, John Common, Chong Yap Seng, Elizabeth Huiwen Tham, Lynette Pei-Chi Shek, Evelyn Xiu Ling Loo, John Chambers, Yik Weng Yew, Marie Loh, Thomas L. Dawson

**Affiliations:** A*STAR Skin Research Labs (A*SRL), Agency for Science, Technology, and Research (A*STAR) & Skin Research Institute of Singapore (SRIS), 11 Mandalay Rd, #17-01, Singapore 308232, Republic of Singapore; DSM-Firmenich, Perfumery and Beauty, Wurmisweg 576, 4303, Kaiseraugst, Switzerland; School of Biological Sciences, Nanyang Technological University, 60 Nanyang Dr, Singapore 637551, Singapore; Genome Institute of Singapore (GIS), Agency for Science, Technology and Research (A*STAR), 60 Biopolis Street, Genome, Singapore 138672, Republic of Singapore; SLING, Singapore Lipidomics Incubator, Life Sciences Institute, National University of Singapore, Singapore 119077, Singapore; Cellivate Technologies, Singapore, 609916; Precision Medicine Translational Research Programme and Department of Biochemistry, Yong Loo Lin School of Medicine, National University of Singapore, Singapore 119077, Singapore; Singapore Institute for Clinical Sciences (SICS), Agency for Science, Technology and Research (A*STAR), Singapore; Department of Paediatrics, Yong Loo Lin School of Medicine, National University of Singapore (NUS), Singapore; Khoo Teck Puat-National University Children’s Medical Institute, National University Health System (NUHS), Singapore; Human Potential Translational Research Programme, Yong Loo Lin School of Medicine, National University of Singapore, Singapore; Lee Kong Chian School of Medicine, Nanyang Technological University, Clinical Sciences Building, Singapore, 308232; Department of Epidemiology and Biostatistics, School of Public Health, Imperial College London, St Mary’s Campus, London, United Kingdom W2 1NY; National Skin Centre, Singapore 308205; Department of Drug Discovery, College of Pharmacy, Medical University of South Carolina, USA

**Keywords:** Lipid mediators, skin microbiome, *Malassezia*, menopause, puberty, sebaceous gland, co-cultures, cytokines

## Abstract

While research into gut-microbe interactions is common and advanced, with multiple defined impacts on human health, studies exploring the significance of skin-microbe interactions remain underrepresented. Skin is the largest human organ, has a vast surface area, and is inhabited by a plethora of microorganisms which metabolise sebaceous lipids. Sebaceous free fatty acids are metabolized into bioactive lipid mediators with immune-modulatory properties by skin-resident microbes, including *Malassezia*. Intriguingly, many of the same lipid mediators are also found on human skin, implying these compounds may have microbial or mixed microbial/human origin. To support this hypothesis, we isolated lipids and microbial DNA from the skin of prepubescent, adult, pre- and post-menopausal volunteers and performed correlational analyses using skin lipidomics and metagenomics to compare lipid mediator profiles and microbiome compositions on skin with either low or high sebaceous gland activity. We found that specific microbial taxonomies were positively and negatively correlated with skin lipid mediator species with high statistical significance. 2D *in vitro* co-cultures with *Malassezia* and keratinocytes also directly linked the production of specific lipid mediators, detected on healthy human skin, to secretion of immuno-stimulatory cytokines. Together, these findings further support the hypothesis that microbial-derived skin lipid mediators influence healthy skin homeostasis and skin disease development and progression, thereby spotlighting the relevance of the skin microbiome’s footprint on human health.

## 1. Introduction

Similar to the immediate area surrounding the plant roots, the rhizosphere, the human/mammalian skin surface supports thriving, complex microbiological activity and is the preferred site for colonization by a large and diverse community of bacteria, fungi, and viruses (1). While many skin host-microbe interactions are commensal or mutualistic, some may cause harm to the host (2).

*Malassezia* yeasts are the most prominent eukaryotic, and normally well tolerated, habitants of human and animal skin. Under certain conditions, however, they become pathogenic and cause multiple common skin disorders such as dandruff, seborrheic dermatitis, and pityriasis versicolor, and are linked to exacerbation of atopic dermatitis and psoriasis (3). Outside of a dermatological context, *Malassezia* is also associated with Parkinson’s disease (4), inflammatory bowel disease (5), accelerated pancreatic oncogenesis (6) and significantly reduced survival rates in breast cancer (7, 8). Over the course of evolutionary adaptation to inhabit the mammalian skin niche, *Malassezia* lost the genes involved in *de novo* lipid synthesis, putatively due to uptake of the rich source of human derived sebaceous skin lipids. Given this lipid-dependency, *Malassezia* is predominantly found on lipid-rich skin regions with high sebaceous gland activity, and as such may modulate lipid metabolic activity on skin (9). Indeed, *Malassezia* has been found to not only feast on epidermal lipids, but also to produce several hundred lipidic compounds *in vitro*, including integral constituents of the skin barrier (10). Besides *Malassezia*, other lipophilic microorganisms such as *Cutibacterium* and *Corynebacterium* species inhabit human skin, thus widening the potential interactive lipid metabolic network and crosstalk between the human host and the microbial community.

The cutaneous barrier protects the human body from harsh environmental conditions by reducing excess transepidermal water loss (TEWL) and preventing entrance of pathogenic microorganisms and damaging chemicals. The stratum corneum epidermal lipid matrix, the skin’s outermost layer, consists of 3 main lipid classes; ceramides, cholesterol, and free fatty acids (FFAs). It is not far-fetched to assume that skin-resident lipophilic microorganisms are capable of modifying the lipid profile on the surface of human skin and as such influence skin barrier function and consequently cutaneous health. Indeed, levels of epidermal long-chain unsaturated FFAs (potential substrates for synthesis of lipid mediators with potent biological activities) and several ceramide subspecies are strongly positively correlated with abundance of *Cutibacteria*, *Corynebacteria* and *Staphylococcus* species, respectively (11, 12). It is important to note that unfortunately these studies only probed bacterial communities and the important skin fungal community was not considered. Indeed, when considering not only the proportion of microbial genomes but including the relative size and biomass, the skin fungal community is surely worth consideration (13). Indeed, the lipophilic species *Cutibacterium acnes* has been implicated in enhanced neutral lipid secretion by sebaceous glands (14), and as such may regulate the lipid composition on the skin surface and thereby enhance its own microbial niche. Whether the lipid composition of the *stratum corneum* determines the microbial profile on skin, or whether the dependency and influence is bidirectional, remains to be further investigated.

Interestingly, correlational analyses between gut microbiome, circulating serum, and tissue lipids are well documented both in mice and humans (15). A dysbiotic gut microbiome has the potential to have far reaching consequences for human health by influencing host lipid metabolism and playing a role in metabolic syndrome (16). However, few studies have been performed highlighting linkages between the skin microbial community and *stratum corneum* lipidome (12). To the authors’ knowledge, no research has yet been made publicly available investigating the causal association between skin-resident microorganisms and the bioactive lipid mediators found on human skin. Previous studies using targeted lipidomics analysis revealed that *in vitro* cultured *Malassezia* yeasts are capable of producing oxylipins and eicosanoids, collectively termed lipid mediators (17). It is noteworthy that the ability for biosynthesis of these lipid mediators is not restricted to *Malassezia* species, as the skin resident fungal species *Candida albicans* and *Candida parapsilosis*, as well as the opportunistic fungal pathogen *Aspergillus fumigatus* may also produce similar lipid mediators (18–20).

Lipid mediators are a diverse class of lipids derived from non-enzymatic and/or enzymatic oxygenation of long-chain polyunsaturated omega-3 or omega-6 FAs (PUFA). The enzymatic reactions are catalysed mainly by lipoxygenases, cyclooxygenases, and epoxygenases of the cytochrome P450 family (21). The PUFA substrates for lipid mediator synthesis are obtained either through *de novo* production following repetitive elongation and desaturation steps or from dietary intake. Consumption of PUFA by skin-resident microorganisms for lipid mediator production is believed to be facilitated by phospholipases, which release PUFA from the *sn-2* position of lipids secreted by host epidermal sebaceous glands. Indeed, the genome of *M. globosa* contains at least 13 potential genes encoding secreted lipases and phospholipases (22). The diverse class of bioactive lipid mediators is characterised by having strong immuno-modulatory functions and being involved in complex host/microbe cross-talk (23). The potent pro-resolving, anti- and pro-inflammatory effects of lipid mediators and their associated receptors have long been known to regulate epidermal cytokine production and secretion (24). For example, several pathogenic fungi, including *Cryptococcus neoformans* and *Candida albicans*, are capable of producing Prostaglandin E2 (PGE_2_), which in turn has been shown to have an anti-inflammatory effect on epithelial cells by modulating IL-8, TNF-α and IL-10 production (19).

Intriguingly, multiple *in vitro* cultured *Malassezia* species produce lipid mediators that are detected on human skin, suggesting these compounds are co-synthesized through metabolic pathways shared between host and microbe, or are potentially of entirely microbial origin (17). Unfortunately, the biosynthetic pathways leading to production of these lipid mediators in *Malassezia* are not yet genomically characterized. These findings are only indicative for a contribution of the cutaneous microbiome to the lipid mediator composition on human skin. To further support the hypothesis that skin lipid mediators with potent immuno-modulatory properties can be of skin-resident microbial origin, this study was designed to reveal significant correlations between skin lipid mediator concentrations and skin microbial community compositions: the sebaceous gland activities in pre-pubescent individuals and post-menopausal women are decreased due to lower hormonal levels, compared to the post-pubescent, and pre-menopausal counterparts, respectively (25). As a consequence, lipid-rich skin regions of adult and pre-menopausal individuals preferentially support colonization by lipophilic skin microbial community components such as *Malassezia* (26), *Cutibacteria,* and *Corynebacteria* (27).

In this study, we perform quantitative and comparative metagenomic and lipidomic analyses of skin surface samples from healthy human subjects and show that different sebaceous gland activities inherently linked to their menopausal and pubescent statuses impact skin microbiome compositions. Importantly, several affected lipophilic microbial community members with attributed PUFA metabolizing capabilities can be significantly associated with divergent skin surface lipid mediator levels. We have also partially recapitulated these findings in a 2D *Malassezia*/keratinocyte co-culture model supporting the relevance of these relationships to human skin health.

## 2. Material and Methods

### 2.1 Materials

All endogenous and isotope labelled lipid mediator standards were purchased from Cayman Chemical (Ann Arbor, MI, USA). Flax paper (Zig-zac, London, UK) and the Cuderm D-Squame tapes (CatLog no: SKU-D100, Clinicalandderm, MA USA) were used for sampling of lipid mediators and DNA, respectively, from human skin.

Keratinocyte serum-free media (K-SFM; Life Technologies, Thermo Fisher Scientific) was used to cultivate N/TERT-1 cells. MILLIPLEX MAP Human Cytokine/Chemokine Magnetic Bead Customised 7-plex Panel (Merck Millipore) was used to quantify cytokine levels.

### 2.2 Sampling of subjects

A total of 100 healthy adult subjects were selected from The Health For Life In Singapore (HELIOS) cohort and constituted of 50 females and 50 males aged 40-71, with different ethnic backgrounds, i.e. 20 South Asian, 60 East Asian, and 20 South East Asian.

A total of 36 healthy pre-pubescent subjects with exclusively East Asian ethnicity were selected from the Growing Up in Singapore Towards healthy Outcomes (GUSTO) cohort and constituted of 18 females and 18 males, all aged 9. Inclusion criteria were normal skin with no obvious skin conditions or disease.

DNA samples were taken from left and right side of cheek by using Cuderm tape strips that were pressed 25-50 times per collection, and either the left or right sample was analysed per subject through DNA sequencing.

Similarly, as previously described (17), skin surface lipid samples from the left and right side of cheek were taken by using an absorbent flax paper disc that was applied to the skin sampling site for 5 minutes using Cuderm tape, and either the left or right sample was analysed using targeted LC-MS/MS. All samples were stored at -80 °C until lipidomics and metagenomics analysis.

### 2.3 Lipidomic and Multiplex cytokine analysis of co-cultures between *Malassezia* species and human immortalized keratinocytes

Human immortalized keratinocytes, N/TERT*-*1 (28) were obtained and cultured in keratinocyte serum-free media (K-SFM) supplemented with 25 µg/ml bovine pituitary extract (BPE), 0.2 ng/ml epidermal growth factor (EGF), and 0.3 mM CaCl_2_, as previously described (29). N/TERT-1 cells were plated in 6 cm tissue culture plates 2 days prior, to reach 80% confluency. Media was refreshed and keratinocytes separately co-cultured for 24 hours at 37 °C (5% CO_2_) with triple-washed *Malassezia* cells (**Figure 4A**) equivalent to 125 µl of exponentially grown *M. globosa* (CBS 7966), *M. furfur* (CBS 7982), *M. sympodialis* (CBS 42132) and *M. restricta* (CBS 7877) cultures. The conditioned and co-cultured media were filtered through 0.2-µm filters and stored at -80 °C prior to experimental analysis. Targeted LC-MS/MS analysis was performed as described below, except that the media was diluted and mixed with an equal volume of extraction buffer (116 mM Na_2_HPO_4_, 42 mM citric acid, pH5.6) prior to SPE.

Luminex analysis was performed to measure the targets IL-10, IL-1α, IL-1β, IL-1Rα, IL-6, IL-8 and TNF-α using the MILLIPLEX MAP Human Cytokine/Chemokine Magnetic Bead Customised 7-plex Panel as previously described (30). Briefly, harvested supernatants and standards were incubated with fluorescent-coded magnetic beads pre-coated with respective antibodies in a black 96-well clear-bottom plate overnight at 4 °C. After incubation, plates were washed 5 times with wash buffer (PBS with 1% BSA and 0.05% Tween-20). Sample-antibody-bead complexes were incubated with Biotinylated detection antibodies for 1 hour and subsequently, Streptavidin-PE was added and incubated for another 30 mins. Plates were washed 5 times again, before sample-antibody-bead complexes were re-suspended in sheath fluid for acquisition on the FLEXMAP® 3D (Luminex) using xPONENT® 4.0 (Luminex) software. Data analysis was done on Bio-Plex Manager^TM^ 6.1.1 (Bio-Rad). Standard curves were generated with a 5-PL (5-parameter logistic) algorithm, reporting values for both mean fluorescence intensity (MFI) and concentration data.

### 2.4 Analysis of lipid mediators using targeted LC-MS/MS

The protocol for lipid mediator extraction from flax paper disks using solid phase extraction (SPE) columns and analysis of lipid mediators by mass spectrometry was performed as described earlier (17), except that the LC/MS-MS system used was based on a reversed-phase chromatographic separation (Analysis column: Phenomenex, Kinetex C8, 2.6 µm,150 x 2.1 mm) on UPLC (Nexera X2 LC-30AD) coupled to a Shimadzu-8060 triple-quadrupole mass spectrometer. The commercially available version 3 of the LC-MS/MS method package “Lipid Mediators” from Shimadzu was used for simultaneous analysis of skin lipid mediators. The method contains analytical conditions, i.e. retention times and MRM transitions, for profiling of 214 lipid mediators, including 18 deuterium-labelled standards. LC-MS/MS parameters for detection of jasmonic acid were applied as described previously (31). Peak identification and integration was performed using the LabSolutions workstation software. Data quality was assessed by calculating the coefficient of variance (CoV) of all analytes in the technical quality control (TQC) pooled lipid extracts, as well as the Pearson’s correlation coefficient (R) to determine the linear response curve of TQC serial dilutions. Peak signal-to-noise ratios were determined using the area under the peaks in samples and blank injections. For data normalization, a single point calibration was used, dividing the peak area of the endogenous analytes by the peak area of internal standards in the same group.

### 2.5 Shotgun Metagenomic Analysis

Genomic DNA was extracted from skin tapes and swabs via a bead beating step on FastPrep-24 Automated Homogeniser (MP Biomedicals), followed by magnetic beads extraction using EZ1 Advanced XL Instrument (Qiagen) with EZ1 DNA Tissue Kit (Qiagen). Firstly, 500 µl of Buffer ATL (Qiagen) was added to each sample and transferred into a Lysing Matrix E tube. Sample tubes were bead-beaten at 4 m/s for 30 seconds twice. After homogenisation, tubes were centrifuged at maximum speed for 5 minutes, with 200 µl of the resulting supernatant transferred into 2 ml EZ1 sample tubes (Qiagen). 12 µl of Proteinase K was added to the supernatant, followed by vortexing and incubation at 56 °C for 15 minutes. Finally, samples were transferred to the EZ1 Advanced XL Instrument (Qiagen) for purification, with a final eluate of 50 µl in buffer EB.

Purified genomic DNA underwent NGS library construction steps using NEBNext® Ultra™ II FS DNA Library Prep Kit according to the manufacturer’s instructions. Samples added with fragmentation reagents were mixed well and placed in a thermocycler for 10 mins, 37 °C, followed by 30 minutes, 65 °C to complete enzymatic fragmentation. Post fragmentation, adaptor-ligation was performed using Illumina-compatible adaptors, diluted 10-fold as per kit’s recommendations before use. Post-ligation, samples are purified using Ampure XP beads in a 7:10 beads-to-sample volume. Unique barcode indexes were added to each purified sample and amplified for 12 cycles under recommended kit conditions to achieve multiplexing within a batch of samples. Finally, each library sample was assessed for quality based on fragment size and concentration using the Agilent D1000 ScreenTape system. Samples passing this quality control step were adjusted to identical concentrations through dilution and volume-adjusted pooling. Paired-end (2×151bp) sequences were generated from the multiplexed sample pool using the Illumina HiSeq X Ten platform, which yielded around 898,525,434 paired reads in total (from three lanes) with an average of 11,980,339 paired reads per library for samples from HELIOS cohort and 536,079,659 paired total reads with an average 6,306,820 paired reads per library for samples from GUSTO Cohort (See attached metadata Excel file). All sequencing was done in the Novogene sequencing facility per standard Illumina sequencing protocols.

Raw reads were processed using the in-house Nextflow pipeline. First, the raw reads were filtered using FastQC (32) (v0.20.0) (default parameters) to remove the low quality bases and adapter sequences. Next, the reads were mapped to human reference hg19 using BWA-MEM (33) (v0.7.17-r1188, default parameters) and SAMtools (34) (v1.7) to remove human reads. These decontaminated reads which were retained were used for the generation of taxonomic profiles using MetaPhlAn2 (35) (v2.7.7, default parameters). *Malassezia* species were identified using Pathoscope (36).

### 2.6 Quantification of hormonal serum levels

Concentrations of luteinizing hormone (LH), follicle-stimulating hormone (FSH) and estradiol (E2) were measured from fasting blood samples, obtained from female HELIOS volunteers, in singletons on the Siemens ADVIA Centaur XPT Immunoassay System (Erlangen, Germany). Daily quality checks were performed, accompanied by calibration every 28, 14 and 21 days for LH, FSH and E2 respectively.

### 2.7 Statistical Analyses

Prior to statistical analyses, zero values in all datasets were classified as nil. To determine if lipid mediators differ significantly between HELIOS and GUSTO cohorts, clustering analysis was performed using Seaborn 0.11.2 library, and the Z-score, or standard deviation from the mean, was computed per lipid mediator and each subject was clustered using and the Bray-Curtis distance with farthest point algorithm (FPA). To determine differences in lipid mediators and the microbiome between cohorts, a one-way multivariate analysis of variance (MANOVA) was performed using Statsmodels 0.13.2 and permutational analysis of variance (PERMANOVA) was performed based on a previously established method (37), respectively. MetaPhlAn 2.7.7 was used to determine microbial relative abundance, while MetaPhlAn2 was used for taxonomic references. To determine overall microbial taxonomic profile, data derived from MetaPhlAn2 were normalised using unweighted UniFrac distance via ChocoPhlAn 3 tree as phylogenetic reference. Principal component analysis (PCoA) analyses were performed using the UniFrac output. Components with more than one eigenvalue were analysed using 10,000 permutations of PERMANOVA with respect to Kaiser’s criterion, and subsequently, PCoA was re-performed in Seaborn 0.11.2 and SKLearn 0.13.2. For cytokine concentration comparisons between keratinocyte co-cultures and single-culture, a repeated measures two-way ANOVA with Geisser-Greenhouse correction and a posthoc Dunnette’s test in GraphPad Prism 9.5.1. The Kendall’s Tau b correlation analysis was performed in SciPy 1.8.0 in order to determine the correlations between keratinocyte cytokines and lipid mediators. The resultant Kendall’s Tau b correlation p-values were further analysed using the two-stage step-up method of Benjamini, Krieger and Yekutieli at a false discovery rate of 5% in GraphPad Prism 9.5.1.

## 3. Results

### 3.1. Skin lipid mediator and microbiome profiles are significantly different between pre-pubescent and adult subjects

Lipidomic analysis of the cheek skin surface of 100 adult and 36 pre-pubescent subjects derived from the HELIOS and GUSTO cohorts, respectively, allowed detection and quantification of a total of 43 different lipid mediator species (**Table S1**). Simultaneous skin metagenomic analysis of adjacent sites allowed identification of 370 individual microbial species (**Table S2**. taxonomic profiles obtained from Metaphlan2 showing the relative abundances of taxa from all the samples used here).

**Table S1. Lipid mediators detected on individuals from the HELIOS and GUSTO cohorts.**

**Table S2. Microbial species and relative abundance detected on individuals from the HELIOS and GUSTO cohorts.**

The current state of research is only indicative that skin-resident microorganisms, such as *Malassezia* fungi, may influence lipid metabolism at the surface of human skin and consequently may modulate the cutaneous immune-response (17). Ideally, a study should be designed to link the production of skin lipid mediators to microbial sub-species levels. However, all human skin is populated by a complex microbial community and with the limitations of currently available tools it would be extremely difficult to dissect the roles of specific species or even strains in lipid mediator metabolism on skin.

Skin sebaceous gland activity is significantly lower in pre-pubescent volunteers than adults and lower in post-versus pre-menopausal women (25). Given this premise, we investigated whether changes in skin lipid mediator profiles can be linked to microbial community diversity and/or abundance distribution, potentially in dependence of varying sebaceous gland activity of healthy, unperturbed skin, using a holistic approach (**Figure 1**).

**Figure 1.**
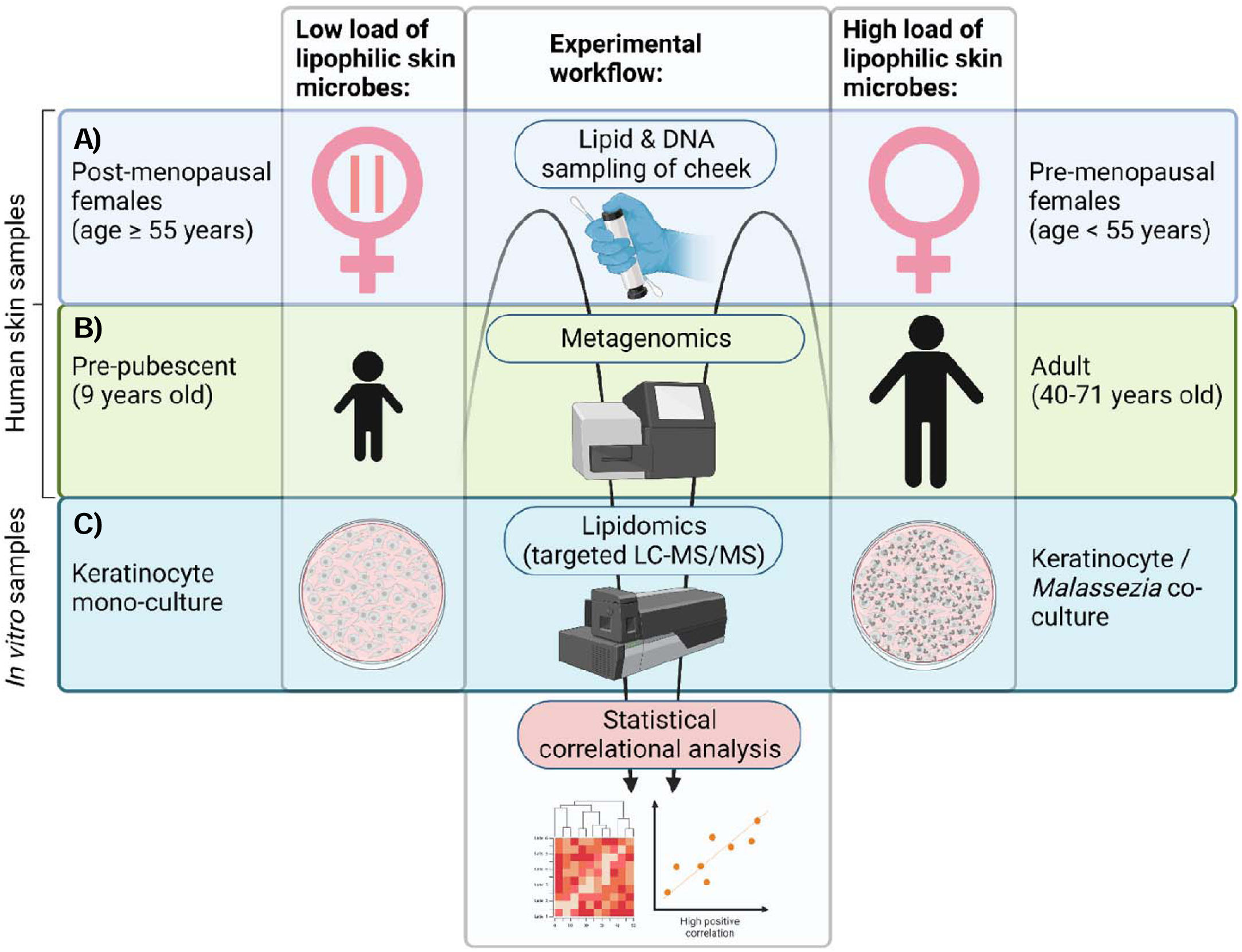
Experimental workflow of targeted lipidomics and metagenomics for correlational analysis of skin lipid mediator profiles and skin microbiome compositions. Cheek DNA and lipid samples were taken from A) post-menopausal and pre-menopausal, B) pre-pubescent and adult volunteers, representing groups with high and low sebaceous gland activities, and consequently high abundance and low abundance of lipophilic skin microbes, respectively. Samples were analysed using targeted LC-MS/MS and metagenomics. C) Supernatants of growth medium from keratinocyte mono-cultures and keratinocyte-*Malassezia* co-cultures, mimicking skin with low and high fungal presence, respectively, were analysed using targeted LC-MS/MS and Multiplex cytokine assay. Correlational analyses were performed for lipidomic/metagenomic, and lipidomic/multiplex data, respectively. Figure created with BioRender.com.

The lipidomic analysis of secreted skin surface metabolites revealed that lipid mediator profiles were drastically different between pre-pubescent (n=36) and adult subjects (n=100): 12 lipid mediator species (out of 43) were significantly increased on skin of the younger cohort, contrasting with 8 species more abundant on the skin of adults (**Figure S1**) following FDR-adjustment. Heatmap clustering and principal coordinate analyses (PCoA) highlighted differences in lipid mediator profiles between skin with low and high sebaceous gland activity, i.e., pre-pubescent and adult skin, respectively (**Figures 2 and 3A**).

**Figure 2.**
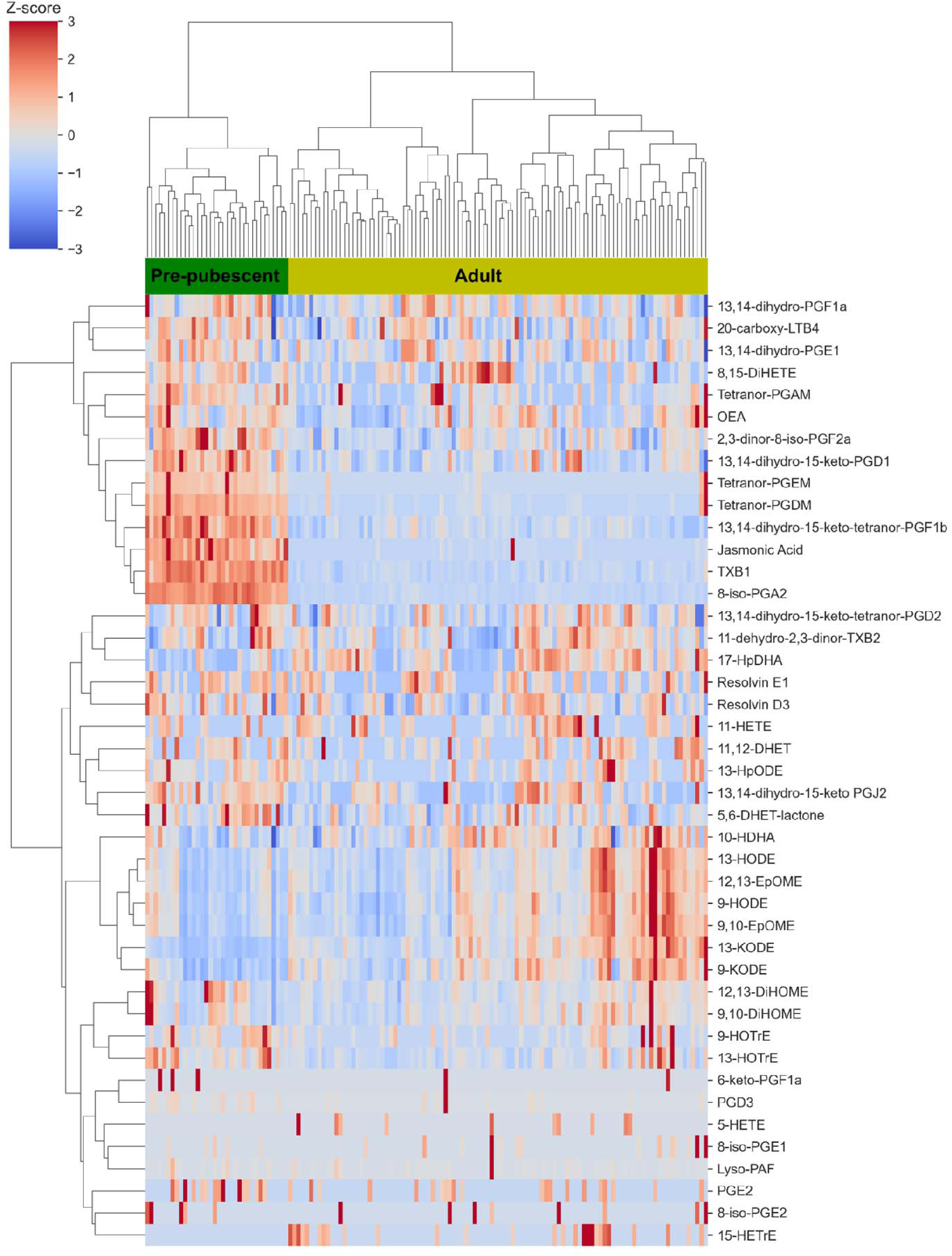
Concentrations of lipid mediators between pre-pubescent and adult skin are different. Skin tape strips from the cheek from the HELIOS and GUSTO cohorts were analysed using targeted mass spectrometry, in which 43 unique lipid mediator species were identified. Red and blue indicates higher and lower lipid concentration relative to the mean, respectively.

**Figure 3.**
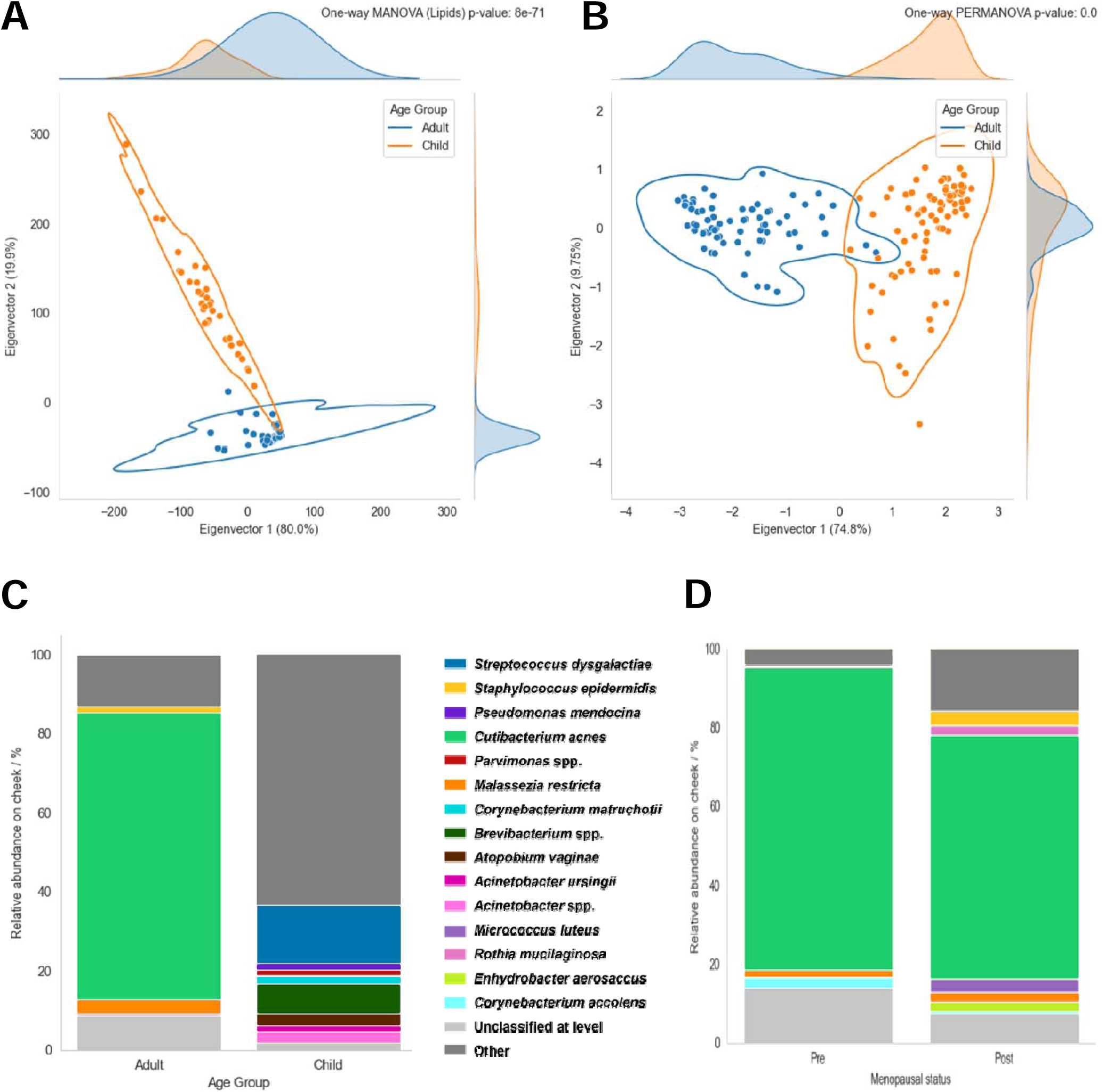
Skin lipid mediator profiles and skin microbiome diversities are different from adult to pre-pubescent, and pre-menopausal to post-menopausal individuals. A) Cohort-level differences in lipid mediators between pre-pubescent (GUSTO) and adults (HELIOS). The PCoA displays raw Euclidean distances multiplied by PCoA eigenvalues and analysed using MANOVA with Wilks’ lambda p-value. Each dot represents an individual (outliers not shown) while lines encompassing dots represent clustering at 95% kernel density estimation using Scott’s method. B) Cohort-level differences in microbiome profile between pre-pubescent and adults. The dataset was analysed using a one-way PERMANOVA with exact p-value. C) Pre-pubescent skin microbiome is more diverse than on adult skin, and D) post-menopausal skin demonstrates more microbiome diversity than pre-menopausal skin. MetaPhlAn2-derived relative abundances are plotted. “Others” refer to microbes that have a count of less than 1.5% while “unclassified at level” refer to microbes at and above the genus level (GUSTO: n = 34; HELIOS: n = 100; [HELIOS] pre-menopausal: n = 25; [HELIOS] post-menopausal: n = 25).

**Figure 4.**
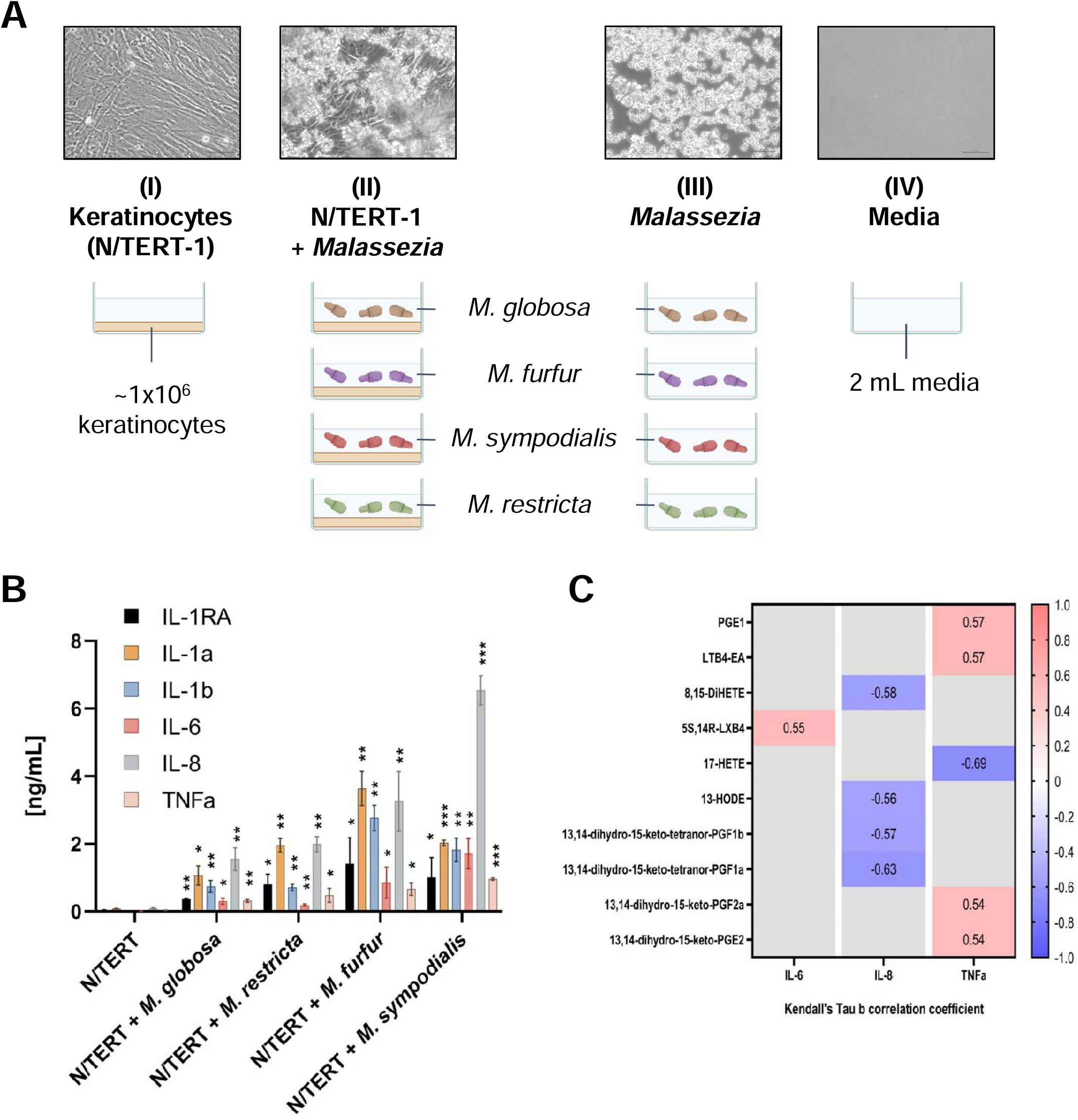
*In vitro Malassezia-*keratinocyte co-cultures associate lipid mediator with cytokine secretion. A) Schematic of the co-culture conditions. Conditioned media from I) mono-cultured human immortalized keratinocytes (N/TERT-1), II) co-cultures with 4 different *Malassezia* spp., III) which were mono-cultured independently, and IV) mock cultured media alone, were used for quantification of secreted cytokines and lipid mediators using cytokine Multiplex and targeted LC-MS/MS analysis, respectively. Representative light microscopic pictures show corresponding mono- and co-culture conditions between N-TERT-1 and *M. globosa* cells. B) Cytokine levels are significantly increased in *Malassezia*-N/TERT-1 co-culture media compared to N/TERT-1 mono-cultures. Bar graphs represents the mean of three biological replicates (n = 3) and error bars represent 95% confidence intervals (CI). Comparison for a specific cytokine was made between the mono- and co-cultures. \**p* < 0.05; \*\**p* < 0.01; \*\*\**p* < 0.001. C) Specific lipid mediator species are correlated with cytokines in *in vitro* culture. Correlation and statistical tests were performed using Kendall’s Tau b with 5% false discovery rate (FDR) correction. Red indicates positive correlation while blue indicates negative correlation. Strength of correlation ± 0.30 or above is classified as strong.

In agreement with previous studies (38), our shotgun metagenomic analysis showed significant differences in microbial communities between pre-pubescent and adult skin (**Figure 3B**). Also, the skin microbiome of the pre-pubescent group was more diverse than the one of the adult group (**Figures S2A and S2B**). Furthermore, as previously shown (27, 39), lipophilic and lipid-dependent microorganisms, such as *Cutibacterium acnes* (formerly *Propionibacterium acnes*) and *Malassezia restricta* were more abundant on adult skin as compared to pre-pubescent skin (**Figure 3C**), likely due to differing age-dependent sebaceous gland activities (25). No statistically significant gender-related differences in lipid mediator levels or skin microbiome profiles were observed (not shown).

Given that cultured *Malassezia* yeasts are capable of producing the same lipid mediators (17) also detected in the course of this study, it was reasoned that the fungal genus’ prevalence on adult skin may be linked to lipid mediator production on the skin epidermis.

### 3.2 Different correlations between the skin microbiome and lipid mediators exist in the pre-pubescent versus the adult cohort

Next, a correlational analysis between skin lipid mediator profiles and skin microbiome at different taxonomic levels was performed to identify and highlight potential engagements of skin-resident microorganisms in epidermal lipid mediator metabolism. Lipid mediator concentrations on adult skin had a wider range of positive correlations to relative abundances of different microbial taxonomies compared to pre-pubescents (**Figure S3**).

At a Spearman’s p-value level, several detected *Malassezia* species showed statistically significant connections with a variety of lipid mediators. Interestingly, on adult skin the most represented *Malassezia* species are *M. globosa* and *M. restricta* (40), and they were tending to be negatively correlated with respective lipid mediators. This suggests that *Malassezia* species differentially contribute to skin lipid mediator profiles. In agreement with the lower abundance of *Malassezia* species on pre-pubescent skin, fewer significant correlations were observed (**Figure S4**).

In summary, changes in cutaneous microbiome diversity and relative abundance between adult and pre-pubescent cohorts significantly correlated with changes in lipid mediator levels. We hypothesize that the lower sebaceous gland activity in skin of pre-pubescent subjects potentially impacted skin microbial community composition and consequently microbial-driven enzymatic production of lipid mediators.

### 3.3 Skin microbiome diversity and lipid mediator profiles are linked to menopausal status

Limited data is available on facial skin microbiome compositional differences with increasing age in female cohorts, and the few existing studies imply hypothetical connections between skin microbiome community distributions and menopausal statuses only (41–46). Given the observed drastic differences in skin microbiome and lipid mediator profiles between pre-pubescent and adult individuals, it was reasoned that the relatively lower sebaceous gland activity present in the aging skin of post-menopausal women (25) may as well differentially dictate its microbiome composition and consequently its epidermal lipid metabolic activity, compared to the pre-menopausal counterpart.

Concomitant with a decrease in endogenous levels of androgens, sebaceous gland activity decreases progressively after menopause (25), potentially leading to dry and itchy skin. In agreement with previous studies (47), serum levels of estradiol (E2) were significantly decreased in our study, and luteinizing hormone (LH) and follicle-stimulating hormone (FSH) levels were significantly increased in post-menopausal (n=25, aged ≥ 55) compared to pre-menopausal subjects (n=25, aged < 55) (**Figure S5**). Notably, the significantly different skin lipid mediator profiles between pre-menopausal and post-menopausal volunteers (**Figure S6**) correlated well with changes in microbiome richness: Similar to pre-pubescent skin, there was higher overall microbial diversity in post-menopausal skin (**Figures 3D** **and S2B**) and a higher number of correlations between lipid mediator species and specific microbial taxonomies compared to pre-menopausal skin (**Figures S7 and S8**). The observed decrease in relative abundance of lipophilic species like *Cutibacterium acnes* (**Figure 3D**) may be associated with a hormone-induced reduction of sebum secretion common in post-menopausal skin, and therefore may be accountable for changes in bioactive lipid mediator profiles.

### 3.4 The Singaporean-Asian ethnicity has a different skin lipid mediator profile but a similar skin microbiome composition

Few studies compared racial differences in sebum composition and the results are controversial: Several studies imply significant differences in sebaceous gland activity and sebaceous lipid profiles between Caucasian, Asian, and American African individuals (48, 49), but another cohort study did not detect significant changes in quantities of sebum lipids using GC-MS analysis of skin tape strips (50). To the authors’ best knowledge, no study has yet compared sebaceous lipid secretion between Asian ethnic subgroups. In the herewith presented study, the analyzed adult cohort consisted of a combination of Asian ethnicities, including South Asian (n=20), East Asian (n=60), and Southeast Asian (n=20). Sebum secretion was higher in the East Asian group (Chinese) relative to the South Asian (Indian) and Southeast Asian (Malay) groups, with significance achieved between the highest (East Asian) and lowest (Southeast Asian) (**Figure S9**). Our hypothesis predicts that differing sebum lipid content should affect skin microbiome compositions and consequently lipid mediator profiles. However, contrary to our expectations, our study found no significant difference in skin microbial alpha- and beta-diversity between these ethnic groups (not shown), consistent with findings from a previous report, which did not detect differences in *Malassezia* species levels between Chinese, Malay, Indian, and Caucasian ethnic backgrounds (51). Nevertheless, we found that several lipid mediator species were significantly different between the Asian subgroups (**Figure S10**), suggesting that other extrinsic and/or intrinsic factors contribute to the divergent lipid mediator profiles of these different Singapore-resident ethnic groups. Particularly, no lipid mediator species were significantly different between South Asian (Indian) and Southeast Asian (Malay), while the East Asian (Chinese) skin lipid mediator profile significantly segregated, in direct parallel to the sebum amount. The total lipid mediator concentration on skin of the Southeast Asian group was also significantly higher than the two other ethnic groups (Kruskal-Wallis p=1e-06).

### 3.5 2D *Malassezia-*keratinocyte co-cultures connect lipid mediator production to cytokine secretion

Since the above correlations do not imply causation, we further investigated the potential of *Malassezia* to produce bioactive lipid mediators in a defined *in vitro* co-culture system with human immortalized keratinocytes (N/TERT-1), the primary cell type of the skin epidermis (28). Skin is exposed to many extrinsic factors, such as topical cosmetics, solar radiation, and air pollution components which can influence skin microbiome composition (52) and lipid mediator profiles (53). To correct for these uncontrollable variables, a complementary *in vitro* experiment was designed to investigate the direct impact of *Malassezia* yeasts on lipid mediator production by using a comparative model system, allowing the mimicking of skin with high and low fungal burden (**Figures 1 and 4A**).

Targeted lipidomic analysis of conditioned growth medium derived from independent co-cultures between N/TERT-1 cells and 4 different *Malassezia* species revealed that the lipid mediator profiles from respective *Malassezia* and keratinocyte mono-cultures were significantly different from *Malassezia*:N/TERT-1 co-cultures (**Figure S11**). Heatmap clustering analysis of lipidomic data revealed multiple lipid mediator species were either enriched or depleted in the respective mono- and co-culture systems (**Figure S12 and S13**).

To investigate a potential role of lipid mediators in directing the relationship of *Malassezia* yeasts with the human host, selected cytokine levels were simultaneously measured in *Malassezia*/keratinocyte co-culture media using a multiplex cytokine assay. In agreement with previous studies (54), all *Malassezia* species induced, to different degrees, secretion of all analysed cytokines except IL-10 (not shown) (**Figure 4B**). As expected, cytokines were not detectable in *Malassezia* monocultures (not shown). The significant correlations between *Malassezia*-induced lipid mediators and keratinocyte-produced cytokines in co-culture are indicative for a potential of these yeasts to modulate the skin immune response triggered by keratinocytes (**Figure 4C**). Interestingly, in contrast to all other analyzed cytokines, IL-8 was exclusively negatively correlated to all lipid mediator species on human skin (**Figure 4C**). The pro-inflammatory cytokine IL-8 is involved in the killing of *Candida albicans* (55), an opportunistic fungal pathogen that colonizes healthy human skin (56), raising the hypothesis that *Malassezia*-derived lipid mediators may repress IL-8 secretion by keratinocytes and thereby antagonize the antifungal effect. Indeed, cultured *Malassezia* species were capable to produce all the negatively correlated lipid mediators: 13-HODE, 8,15-DiHETE, 13,14-dhk-tetranor-PGF1β, 13,14-dhk-tetranor-PGF1α (**Figure S12**).

Lipid mediators, as well as cytokine levels, were either below the detection limit or significantly lower in raw growth medium and N/TERT-1 monocultures than in the experimental culture conditions. Furthermore, co-culture conditions were well tolerated by *Malassezia* species and the keratinocytes, since the yeasts recovered well on agar growth plates following exposure to 37 °C (5% CO_2_) for 24 hours, and a lactate dehydrogenase (LDH) assay revealed no significant cytotoxicity levels, respectively (not shown).

## 4. Discussion

Several fungal species, including skin resident *Malassezia* yeasts, are capable of producing lipid mediators found on human skin, raising the hypothesis that these compounds may be synthesized either through shared metabolic pathways or be entirely of microbial origin (17). Ideally, an experimental approach would allow perturbance of the skin microbial community with species- or strain-specific antimicrobials for controlled removal to investigate the direct impact of skin microbial species or strains on skin lipid mediator metabolism. However, given the complexity of such a study and the unavailability of such targeted materials, it was instead decided to perform a comparative analysis of lipid mediator profiles and microbiome compositions on healthy, unperturbed human skin with either low or high sebaceous gland activity and hence different microbial communities (**Figure 1**).

We found that in healthy human subjects with differing skin sebum content the microbial communities, as expected, were different. There was higher microbial diversity associated with lower sebum content, both in pre-pubescent volunteers and post-menopausal women. This correlated closely with changes in the skin lipid mediator profiles similarly in all groups. It would be of clinical significance to reveal whether particular microbial lipid metabolism pathways can be fitted to these changes. Future research into genomic characterization of lipid mediator biosynthetic pathways in skin-resident microbes will ultimately allow the identification of targets for clinical applications for skin disease prevention. Further support for the hypothesis that lipid mediator profiles are shaped by differing sebaceous gland activities was found when comparing different Singaporean ethnicities. The East Asian (Chinese) cohort was found to have higher lipid secretion than Southeast (Malay) or South Asian (Indian) subjects, with corresponding changes in lipid mediator richness. It is likely that the factors impacting lipid mediator profiles between ethnicities are far more complex than merely human genetics, sebum secretion, and the microbial community. While the lack of these specific study participants’ metadata hinders the interpretation of the results, multiple theoretical factors could be responsible for these differences: Differing skin care routines, dietary regimens, or genetic differences in skin composition could be accountable for the observed skin microbiome-independent variations in lipid mediator profiles. Indeed, South Asian (Indian) skin and hair care traditionally involves topical application of natural oils derived from olives, sunflower seeds, coconuts and oats (57, 58), which are all rich in essential omega-3 and omega-6 fatty acids, including PUFAs that can impact lipid mediator metabolic pathways on skin. Moreover, mediated via the gut-skin axis, differing dietary habits can have a substantial impact on skin mediator lipidomes. For example, dietary supplementation of omega-3 PUFA, enriched in e.g. flaxseed oil, a commonly used ingredient in Indian cuisine, reduced the production of arachidonic acid-derived lipids with pro-inflammatory properties in the epidermis (59). Last but not least, skin microbiome profiling with higher resolution genome assembly and sequencing depth to enable improved strain-level discrimination could potentially reveal differences in microbiome community distributions at strain levels between different ethnic groups. Alternatively, determination of absolute quantities of skin microbial species could potentially reveal differences in abundance distributions between examined groups, a feature that is currently not feasibly achieved by shotgun metagenomic sequencing.

In support of the hypothesis that sebaceous gland-derived lipids dictate skin microbial richness and composition, which as a consequence modulates skin microbial activity and in turn bioactive lipid mediator production, we found a high number of lipid mediator species either positively or negatively correlated with specific skin microbial taxonomies with high statistical significance. Alternatively, increased sebaceous gland activity could simply increase PUFA substrate availability for lipid mediator production without any impact on skin microbial community composition, explaining the observed differences in lipid mediator profiles. However, in this study total lipid mediator concentrations were significantly higher on pre-pubescent skin with lower sebaceous gland activity (6-fold; p=2e-09) relative to adult skin, and no statistically significant differences in total lipid mediator concentrations were observed between higher sebaceous activity on pre-relative to post-menopausal skin (not shown).

To complement the human study with an *in vitro* approach we co-cultured keratinocytes with varying concentrations of *Malassezia* cells, thereby mimicking skin with low or high fungal presence, respectively. The doses were chosen to bracket presumed concentrations of yeast cells on human skin. We then quantified secreted lipid mediator and cytokine levels. Interestingly, 37% of all quantified lipid mediator species that were either positively or negatively correlated with specific skin microbiome taxonomies in the *in vivo* study overlapped the *in vitro* experiments (**Figure S14**). Consistent with the human study, 33% of lipid mediator species, whose concentrations were at least log 1.5-fold different between *Malassezia* mono- and co-cultures (**Figure S12**), albeit not significantly different from controls based on statistical tests, were also correlated with skin microbiome taxonomies (including Resolvin D3, 8-iso-PGA_2_, 8-iso-PGE_2_, 8-iso-PGE_1_, PGE_2_, 9,10-DiHOME, 12,13-DiHOME, 12,13-EpOME, 9-HODE, 13-HODE, 11,12-DHET, 8,15-DiHETE, 13,14-dhk-tetranor-PGF1b, and TXB1). It should be noted that while the applied corrections for multiple comparisons (i.e. FDR) removed statistical significance, the observed differences may nevertheless be of biological relevance. Therefore, the presented results suggest that the *in vitro* culture model reflects at least to some extent real-life physiological conditions. While initially appearing contradictory, several lipid mediator species previously shown to be produced by cultured *Malassezia* species (17) were negatively correlated with *Malassezia* relative abundances. However, it is noteworthy that changes in relative DNA abundances, as revealed by shotgun metagenomic approaches, do not correlate with absolute microbial abundances. Therefore, we cannot exclude the possibility that a decrease in the proportional abundance of a specific skin microbial species in fact manifests as a change in actual microbial load. Furthermore, the highly complex skin microbial community involves inter- and intra-species signalling and biochemical activities with synergistic and/or antagonistic effects on skin lipid mediator production. Hence, not surprisingly, a direct translation of findings retrieved from highly controllable *in vitro* culture systems to physiological environments is not straightforward.

In this study, we show that *Malassezia* monocultures secreted high quantities of lipid mediator species including 9-HpODE, 9-HODE, 13-HODE, 9,10-EpOME, 9,10-DiHOME, 12, 13-DiHOME, Resolvin D4, 8,15-DiHETE and 10,17-DiHDHA, which were depleted in co-culture systems (**Figure S12**). It is hypothesized that microbial-derived production of certain intermediate and/or end product lipid mediator species on skin may trigger feedback mechanisms leading to their accelerated absorption, biosynthetic completion, and/or degradation by skin epidermal cells and/or other members of the skin microbial community, thus giving a plausible explanation for why certain lipid mediators produced by a specific microorganism may be negatively correlated with this very species under physiological conditions. For example, this study revealed that the 9,10-EpOME precursor, even though produced by *Malassezia in vitro*, was negatively associated with *M. globosa* and *M. restricta* on adult, pre-menopausal and pre-pubescent skin, respectively, whereas the 9,10-DiHOME product was positively correlated with *M. restricta* on adult skin and *M. furfur* on post-menopausal skin. It can be reasoned that the conversion from the 9,10-EpOME precursor diol to its 9,10-DiHOME product may be performed by the soluble epoxide hydrolase (sEH)-mediated (60) rate-limiting step, potentially effectuated by *Malassezia* on skin.

Interestingly, the linolenic acid-derived phyto oxylipin Jasmonic acid (JA) and the endocannabinoid-like compound Oleoylethanolamide (OEA) were amongst the lipid mediator species increased on pre-pubescent skin, were significantly correlated with several skin microbiome taxonomies, and are produced *in vitro* by cultured *Malassezia* (not shown). Given that endocannabinoid lipid mediators have been implicated in regulatory functions in skin health and disease (61), an interference with the skin endocannabinoid system by skin microbes is plausibly involved in healthy skin homeostasis. Why *Malassezia* produce JA on human skin is a mystery but given that JAs regulate plant immunomodulatory functions, it is hypothesized they similarly influence human skin immunology and hence health and disease. Furthermore, it is possible that *Malassezia* retained JA production as they are evolutionarily derived from plant pathogens like *Ustilago* (62).

Psoriasis is a common, long-term immune-mediated skin disease that has been associated with *Malassezia* (3). Several lipid mediators, such as 8-/12-/15-HETE, 9-/13-HODE, 9-/13-KODE and 9,10-EpOME were shown to be significantly increased in psoriatic skin of adults with a mean age of 48 (63). This study showed that of those, 9-/13-HODE, 9-/13-KODE and 9,10-EpOME were significantly elevated in adult skin compared to pre-pubescent skin. Consistent with the finding that *Malassezia restricta* is proportionally less represented on pre-pubescent skin, and the fact that psoriasis is more common in post-pubescent patients (64), this study further supports the hypothesis that *Malassezia-*derived lipid mediators may be involved in modulating the human skin immune response and may influence skin disease development and progression, thereby putting a spotlight on the relevance of the skin microbiome’s footprint on human health.

Also of note, PGE_2_ and PGD_2_ have been implicated in hair growth stimulation and inhibition, respectively (65) and their concentrations were increased in co-culture conditions. Given the proximity of the sebaceous gland, being a part of the human hair follicle, to the habitat of lipid-dependent *Malassezia* yeasts, a direct role of this fungus in scalp health and hair growth should be considered in future studies.

## 5. Conclusions

We have shown that in periods of life change where skin sebaceous lipid levels are altered, namely puberty and menopause, specific members of the skin microbial community can be linked with high statistical significance to changes in skin lipid mediator species. We have also shown that in an *in vitro* model of *Malassezia*/keratinocyte interactions, multiple lipid mediator species are impacted in a similar manner to those correlations on healthy human skin. While this study only shows a correlational interaction between skin microbiome and lipid mediator levels, the presented results imply that there is a complex microbe/host communication system present on healthy human skin, and that skin health and disease may be influenced by the skin microbial community through polyunsaturated lipid mediators. This should encourage further investigations into potentially more directly causative links.

## 6. Declarations

### Ethics approval and consent to participate

All volunteers from the HELIOS and GUSTO studies provided written informed consent prior to study commencement and were sampled in accordance with the Declaration of Helsinki, and both studies were approved by the Nanyang Technological University (NTU) Institutional Review Board (IRB-2016-11-030), the National Healthcare Group Domain Specific Review Board (D/09/021) and SingHealth Centralised Institutional Review Board (CIRB 2018/2767; 2009/280/D), respectively.

### Consent for publication

Consent was sought from study volunteers for the publication of their de-identified processed data in strict adherence to the approved HELIOS (IRB-2016-11-030) and GUSTO (CIRB 2018/2767; 2009/280/D) study protocols.

### Availability of data and material

All data generated or analysed during this study are included in this published article [and its supplementary information files].

## Acknowledgements

Bo Burla and Ji Shanshan for critically evaluating lipidomics data.

---

**Supplemental Figure 1.**
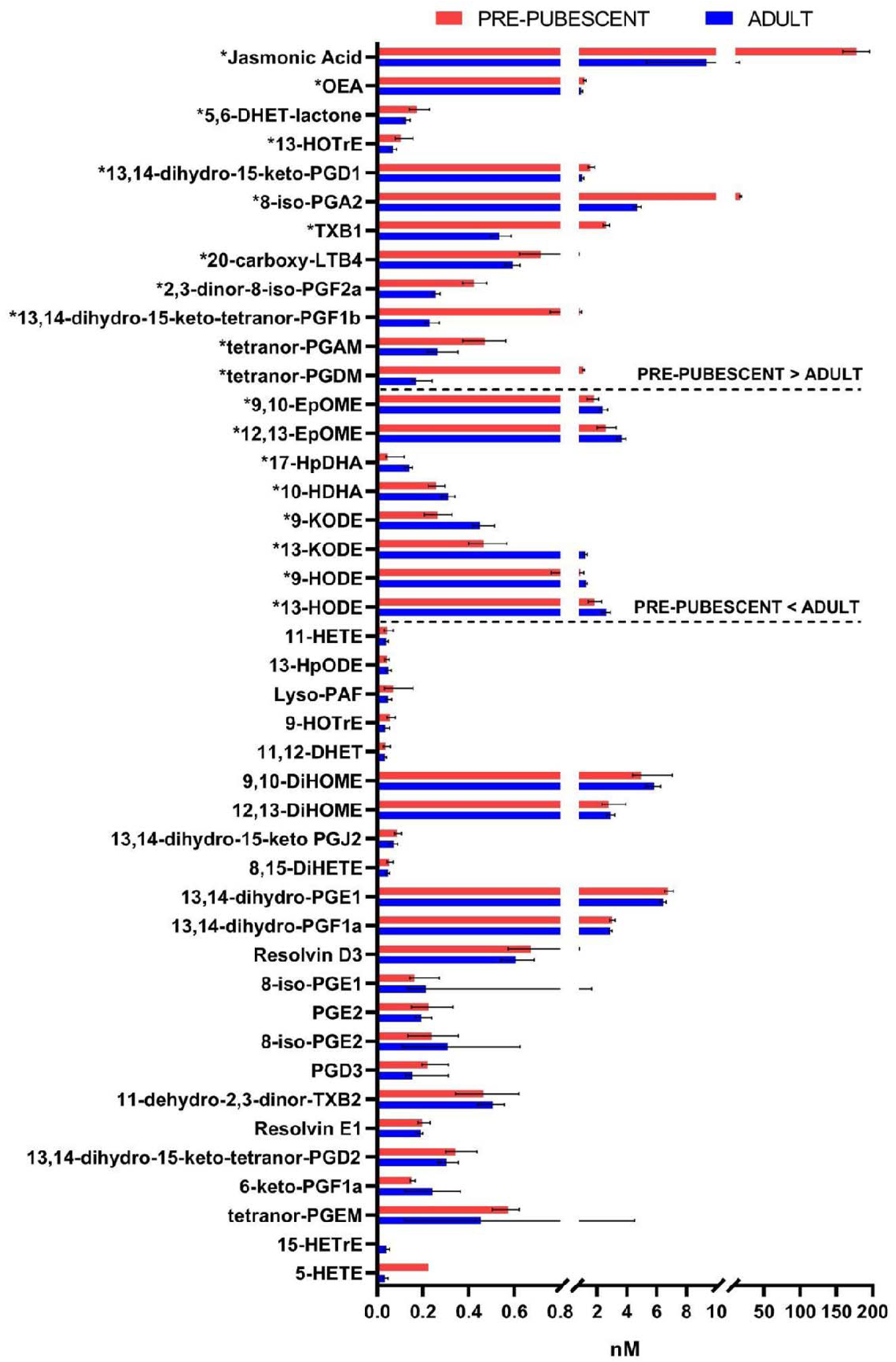
Pre-pubescent and adult cheeks have different lipid mediator concentrations. Lipidomic analysis was performed on pre-pubescent (GUSTO; n = 36) and adult (HELIOS; n = 100) cheek samples. A total of 43 lipid mediators were detected and the dataset was analysed using multiple Mann-Whitney U tests with 1% FDR correction. A total of 20 lipid mediators were identified as having significantly different concentrations between the GUSTO and HELIOS cohorts. Bar graphs represents the median while error bars indicate the 95% CI. *FDR-corrected discoveries.

**Supplemental Figure 2.**
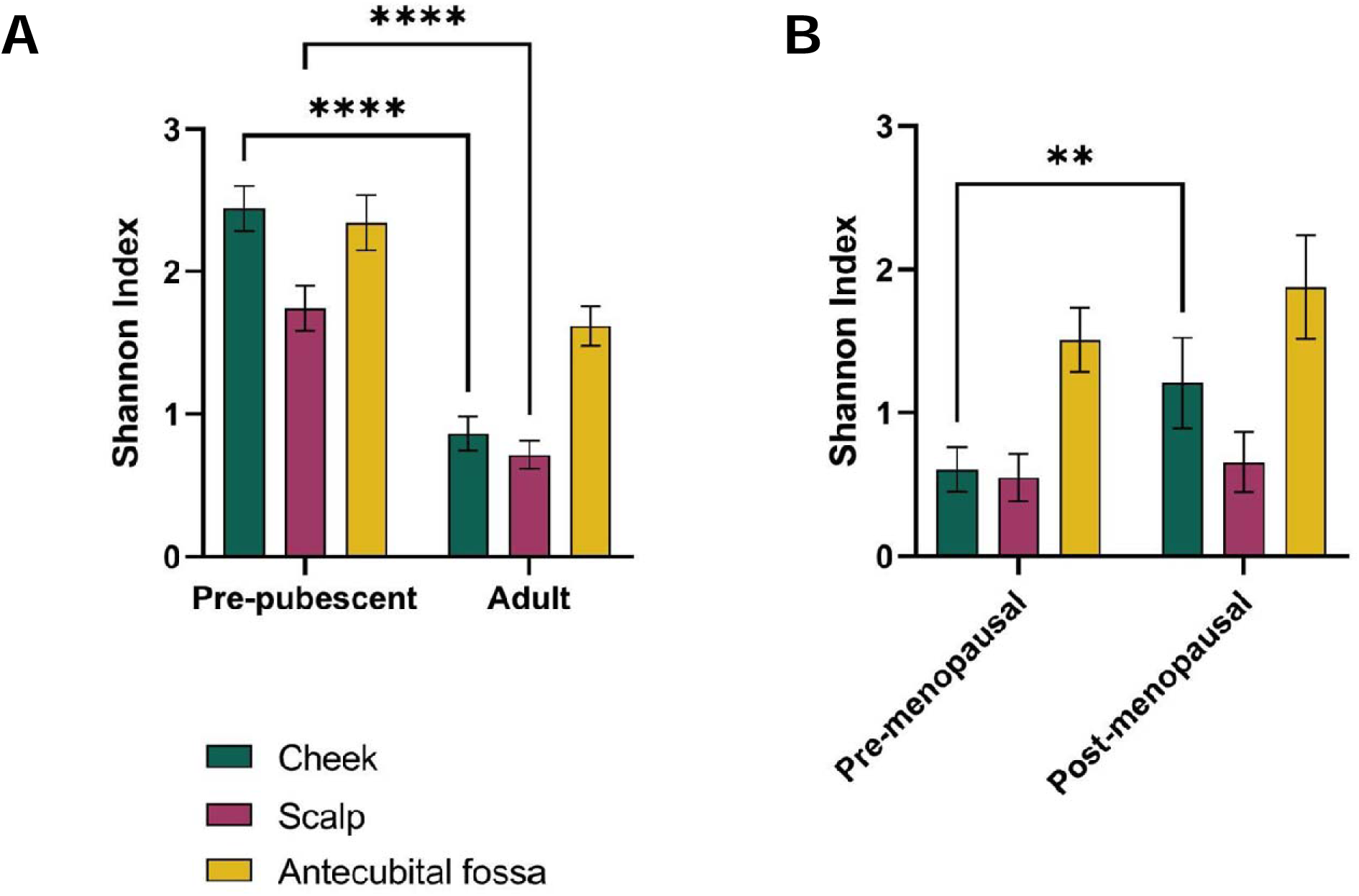
Species diversity is greater in pre-pubescent and post-menopausal groups compared to adult and post-menopausal groups, respectively. Shannon index between A) pre-pubescent (n = 85) and adults (n = 100) and between B) pre- (n = 10) and post-menopausal (n = 25) women. Bar graphs represents the mean, while error bars indicates the 95% CI. Data was analysed with two-way ANOVA with posthoc Tukey’s test. Only significant comparisons are indicated on the graph. \*\**p*< 0.01, \*\*\*\**p* < 0.0001.

**Supplemental Figure 3.**
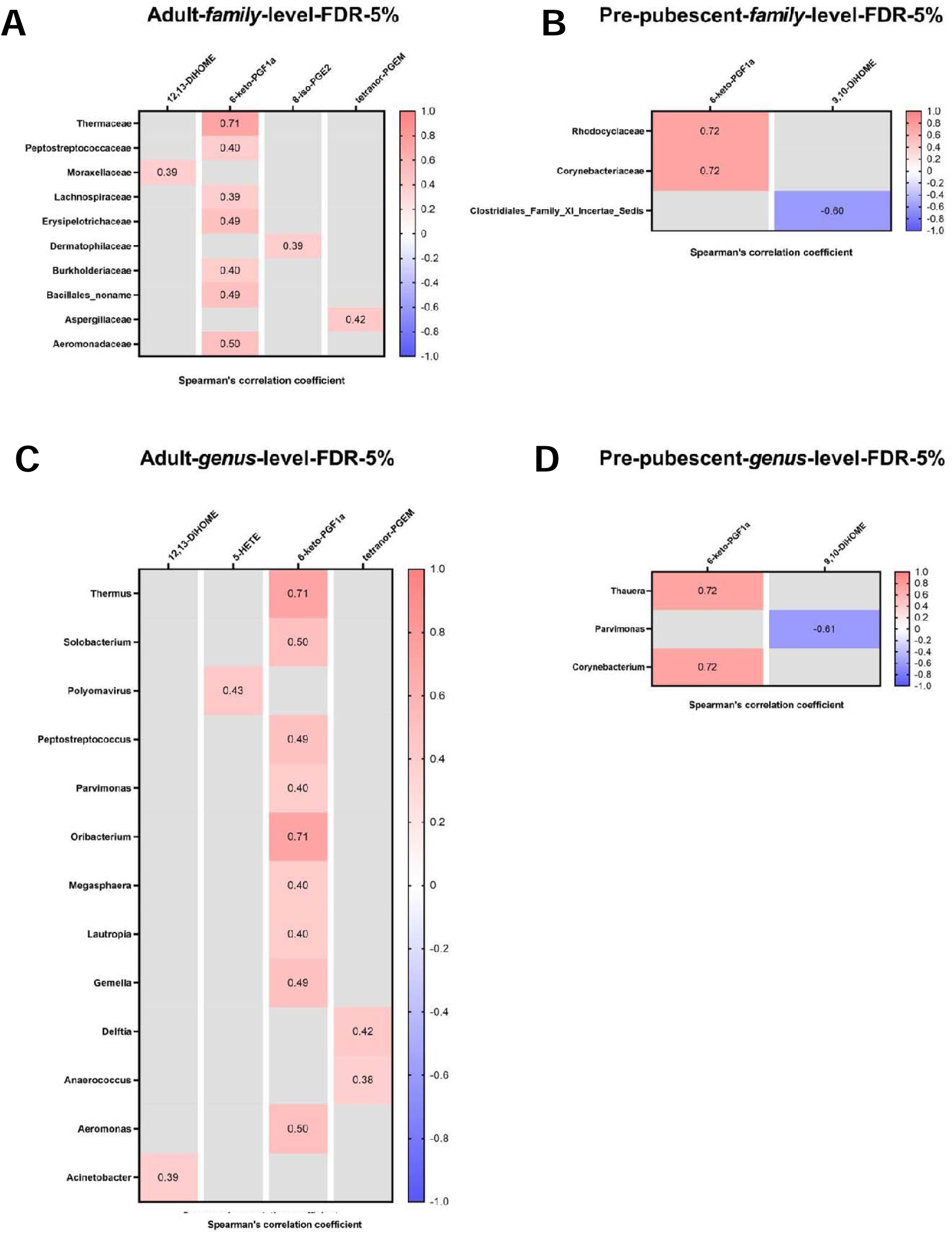
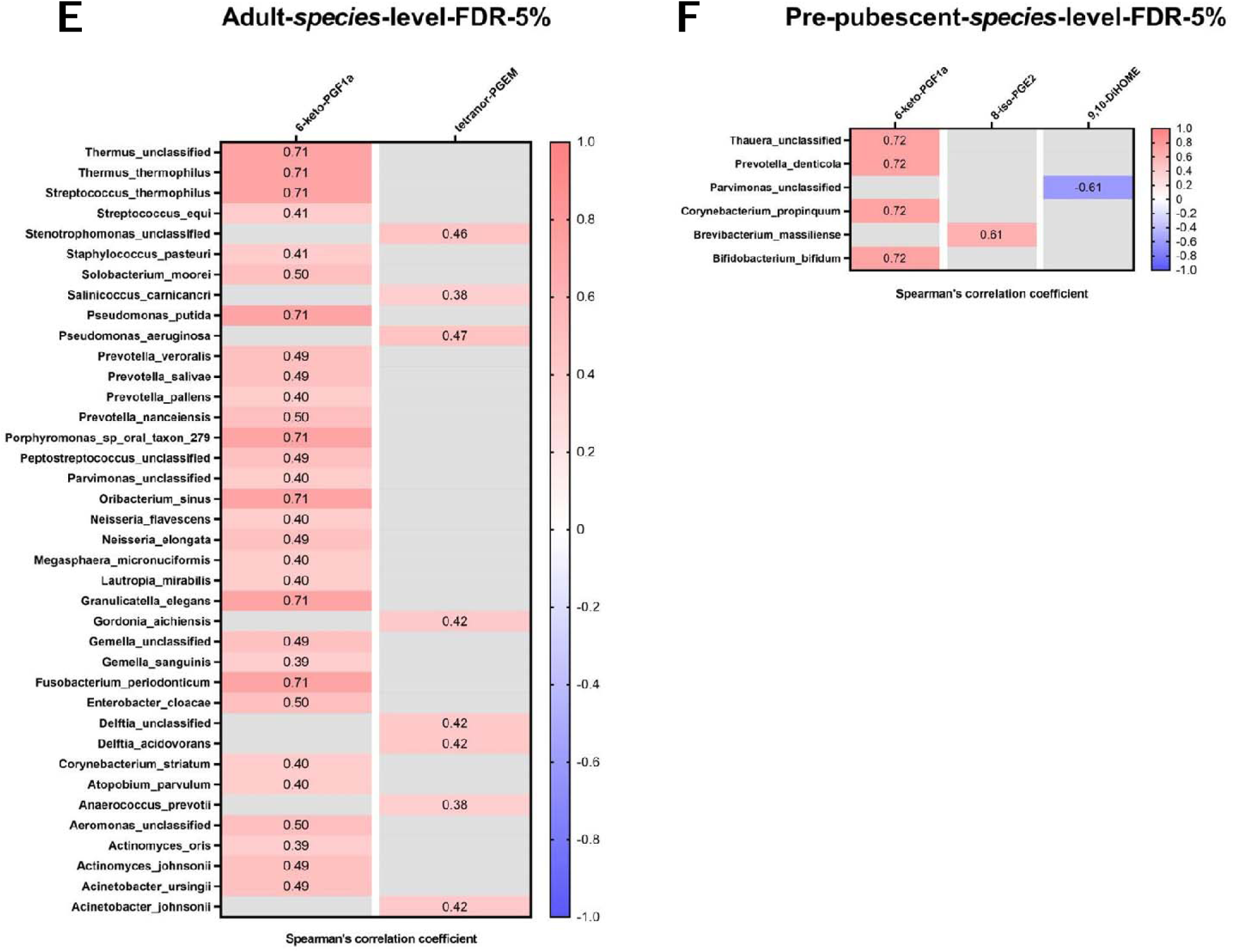
Family-, genus- and species-level correlation of microbes with lipid mediators on pre-pubescent and adult cheeks. Family-level correlations in A) adults and B) pre-pubescent participants. Genus-level correlations in C) adults and D) pre-pubescent participants. Species-level correlations in E) adults and F) pre-pubescent participants. Spearman’s correlation was performed with 5% FDR correction. Only FDR-corrected discoveries are plotted. Red indicates a positive correlation while blue indicates a negative correlation. The strength of correlation is categorized as; ± 0 to 0.09: negligible, ± 0.10 to 0.39: weak, ± 0.40 to 0.69: moderate, ± 0.70 to 0.89: strong, and ± 0.9 to 1: very strong. A perfect positive correlation is +1, while a perfect negative correlation is -1.

**Supplemental Figure 4.**
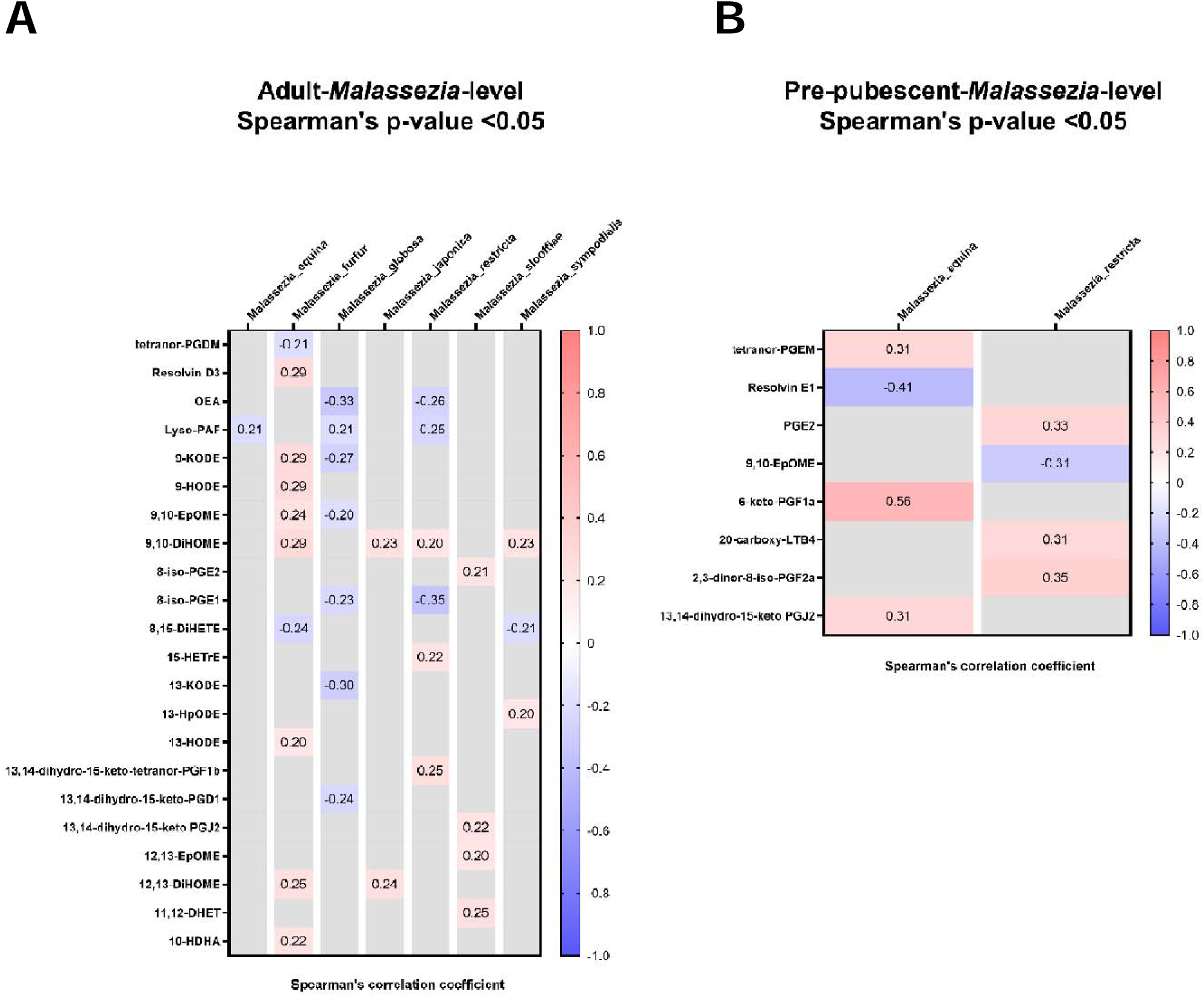
Correlation of lipid mediators with the presence of *Malassezia* species on pre-pubescent and adult cheeks. Correlations in A) adults and B) pre-pubescent participants. Spearman’s correlation was performed with 5% FDR correction. No discoveries were found after FDR correction. Therefore, data plotted were from correlations that had a Spearman’s p-value of < 0.05. Red indicates a positive correlation while blue indicates a negative correlation. Categorization of correlation strength is found in figure legend of S3.

**Supplemental Figure 5.**
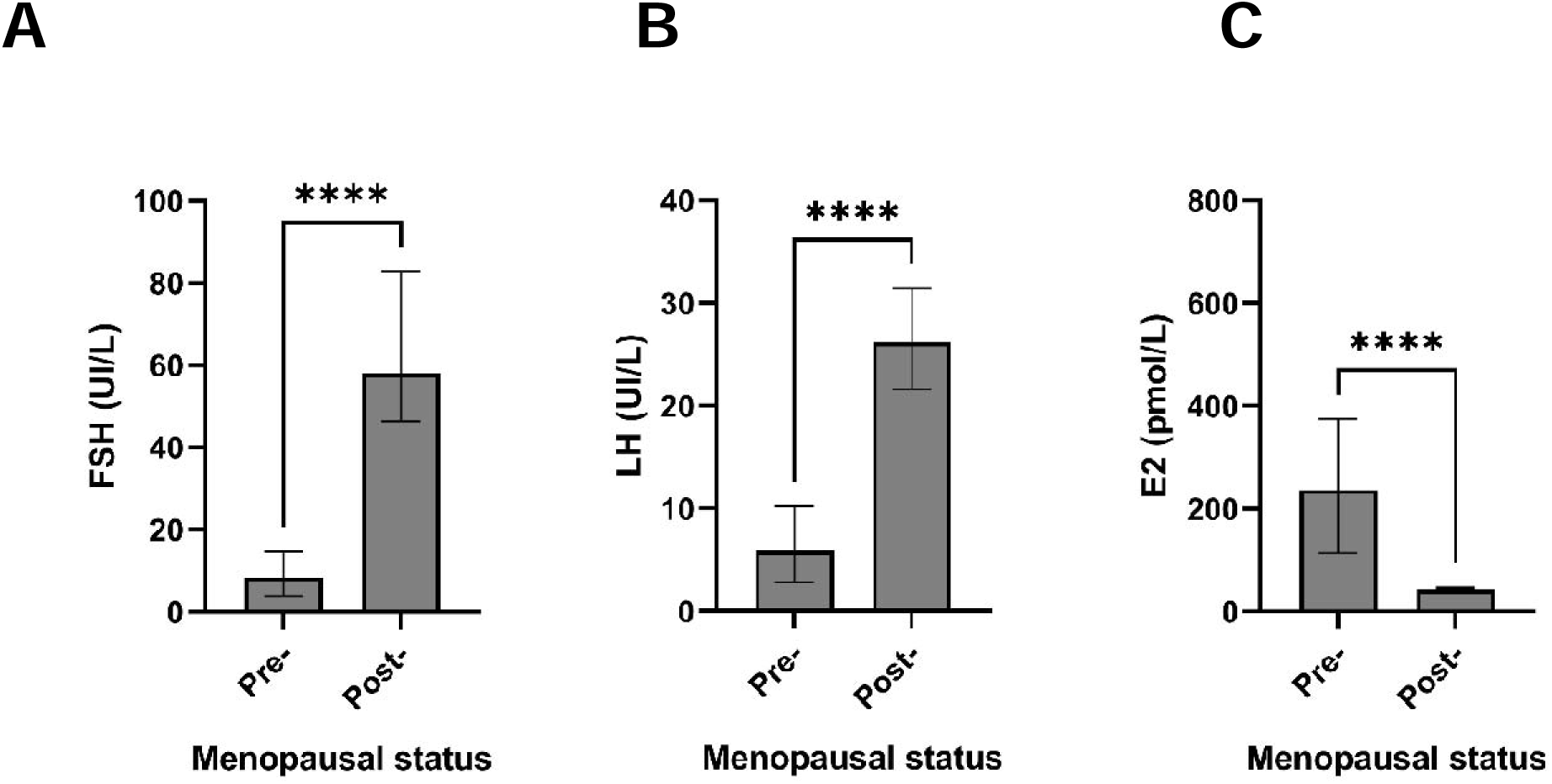
Pre- and post-menopausal women have differing serum hormone concentrations. A) Follicular stimulation hormone (FSH), B) luteinizing hormone (LH), and C) estradiol hormone (E2) concentration. Bar graphs represents the medial hormone concentration of 25 individuals in each group (n = 25). Error bars indicate the 95% CI. Data was analysed using the Mann-Whitney U test. \*\*\*\**p* < 0.0001.

**Supplemental Figure 6.**
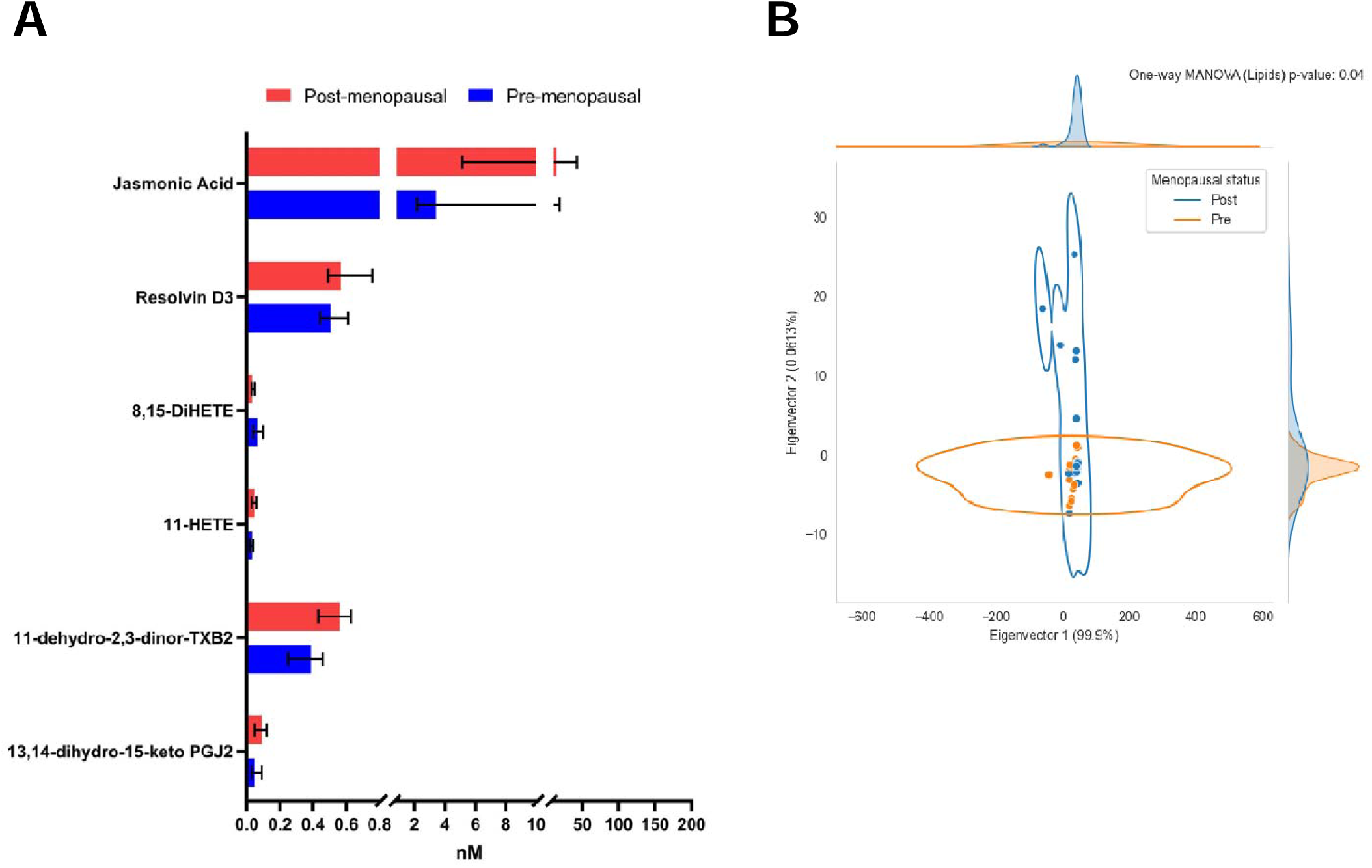
Cheek lipid mediator concentrations are affected by pre- and post-menopausal statuses. A) Concentrations of lipid mediators identified as different between pre- and post-menopausal women. Bar graphs represent the median lipid concentration on the cheeks of pre- (n = 25) and post-menopausal (n = 25) women. Error bars indicate 95% CI. Data was analysed using multiple Mann-Whitney U tests with 5% FDR correction. No discoveries were made using the 5% FDR correction. Therefore, significant differences based on Mann-Whitney U test without FDR correction have been plotted. B) The PCoA plot displays raw Euclidean distances of lipid concentrations multiplied by PCoA eigenvalues. Dots represent individual data points. One-way MANOVA was then performed, with the Wilks’ lambda p-value indicated in the graph. Outliers are not shown in the graph but were used in the statistical calculations. Lines encompassing dots represent clustering at 95% kernel density estimation using Scott’s method.

**Supplemental Figure 7.**
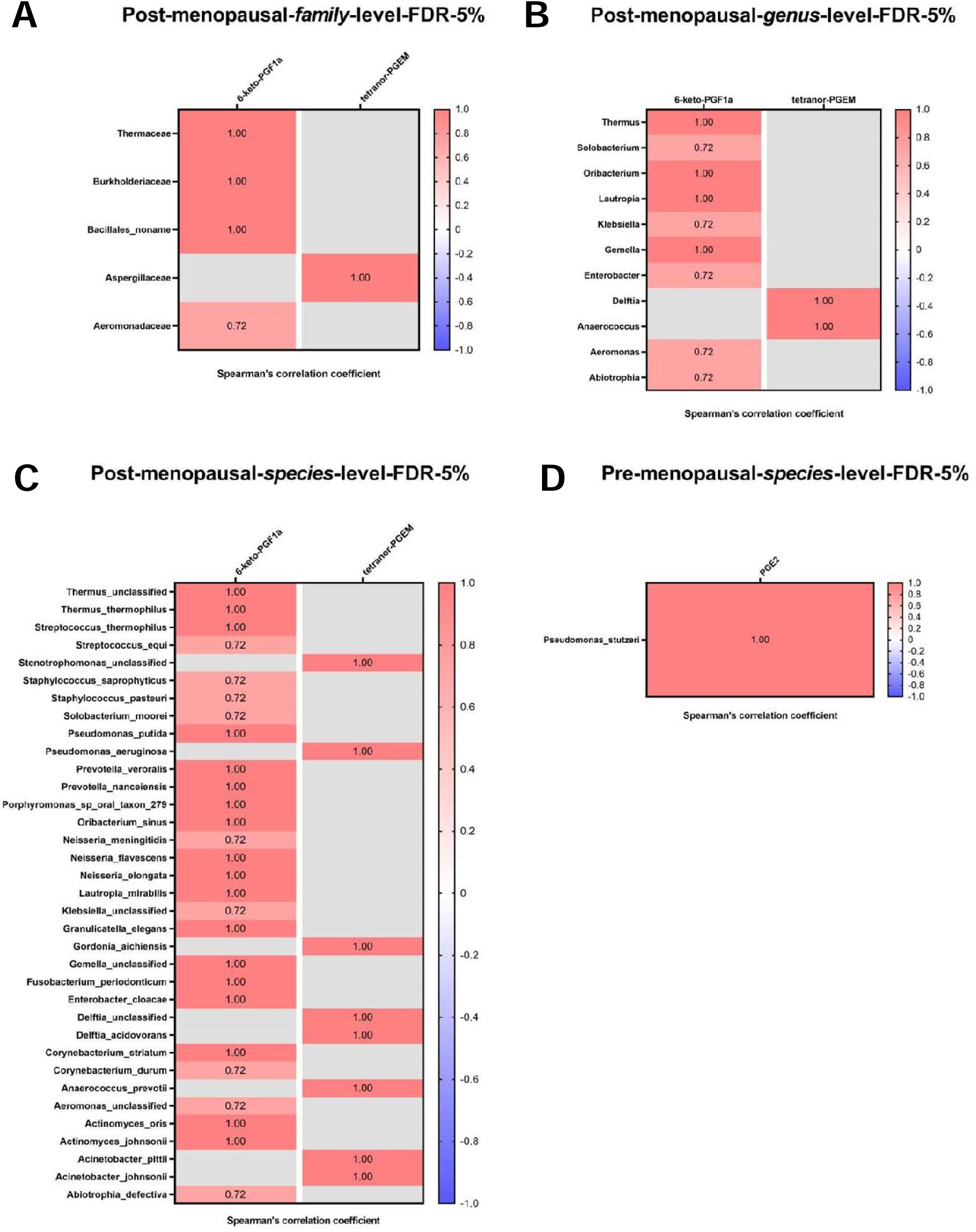
Family-, genus- and species-level correlation of microbes with lipid mediators on pre- and post-menopausal women cheeks. A) Family-, B) genus-, and C) species-level correlations in post-menopausal women. D) Species-level correlations in pre-menopausal women. Spearman’s correlation was performed with 5% FDR correction. Only FDR-corrected discoveries are plotted. Red indicates a positive correlation while blue indicates a negative correlation. Categorization of correlation strength is found in figure legend of S3. No correlation discoveries were made between lipids and microbiome for pre-menopausal women at family- and genus-level.

**Supplemental Figure 8.**
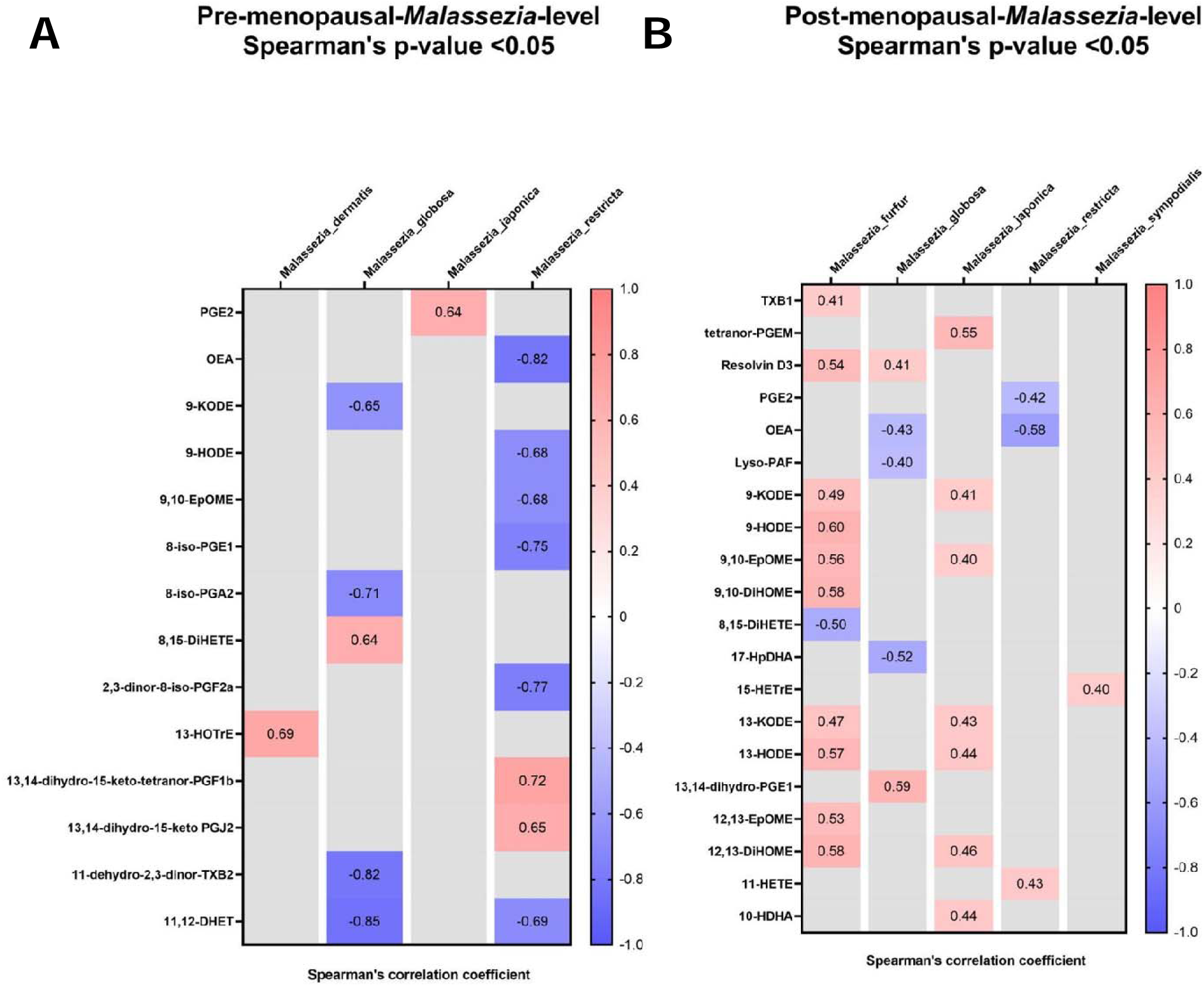
Correlation of lipid mediators with the presence of *Malassezia* species on pre- and post-menopausal women cheeks. Correlations in A) pre- and B) post-menopausal women. Spearman’s correlation was performed with 5% FDR correction. No discoveries were found after FDR correction. Therefore, data plotted were from correlations that had a Spearman’s p-value of < 0.05. Red indicates a positive correlation while blue indicates a negative correlation. Categorization of correlation strength is found in figure legend of S3.

**Supplemental Figure 9.**
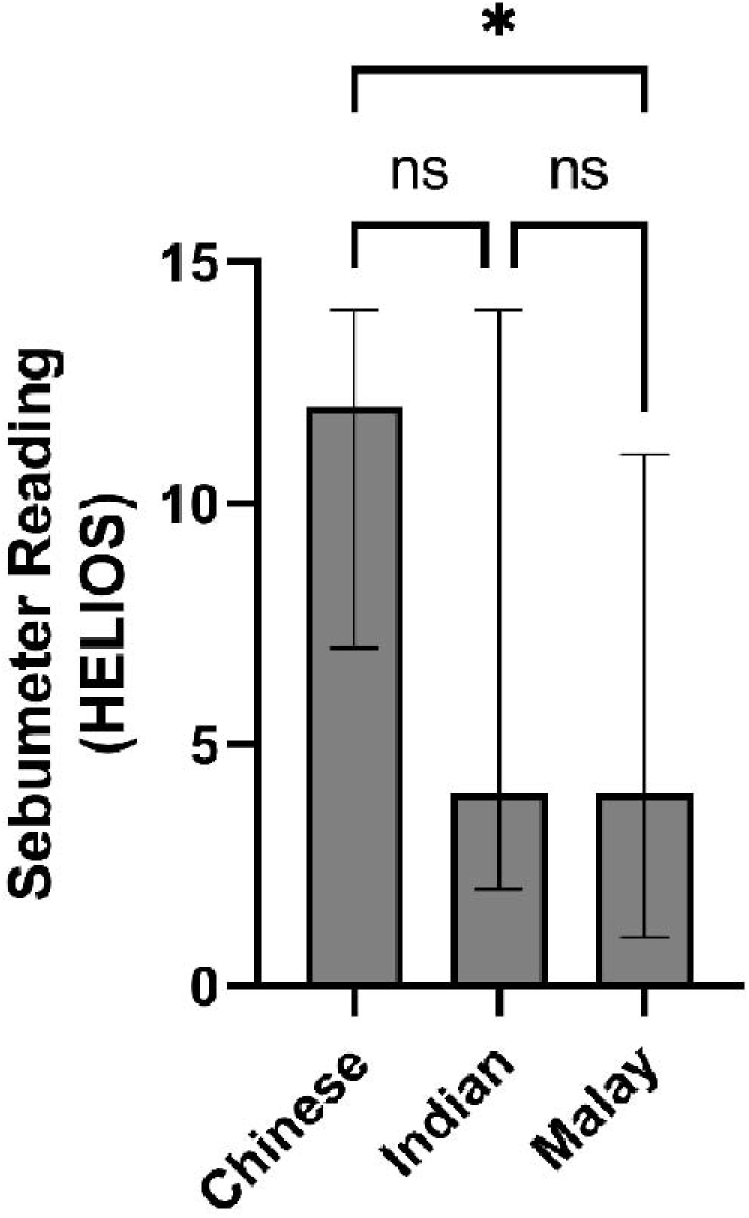
Sebumeter readings are different between East Asian (Chinese) and Southeast Asian (Malay) groups. Bar charts represent the median of groups with 95% CI. Data was analysed using Kruskal-Wallis with posthoc Dunn’s test. \**p* < 0.05.

**Supplemental Figure 10.**
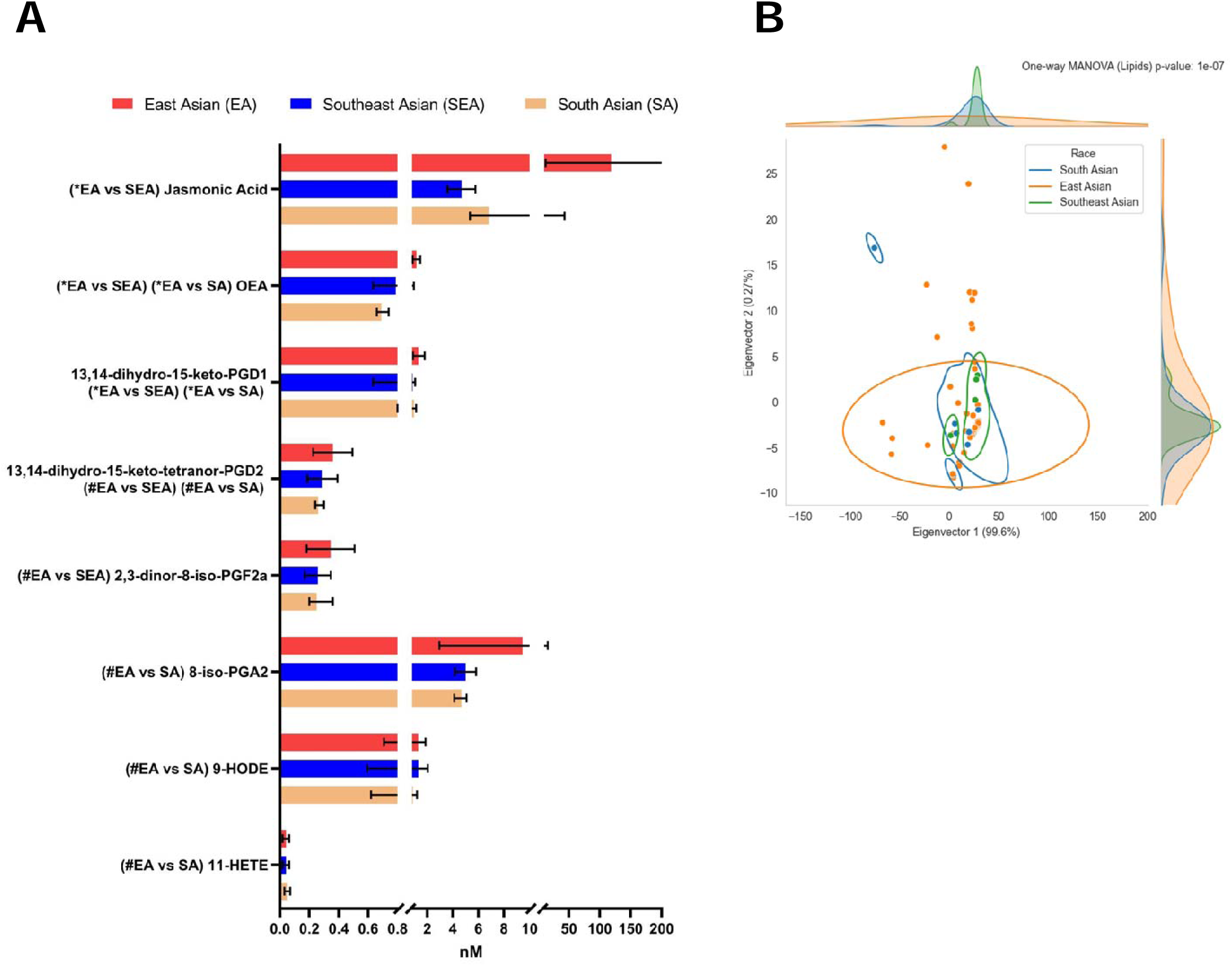
Cheek lipid mediator concentrations are affected by ethnicity. A) Median lipid concentrations between ethnic groups. Error bars indicate 95% CI. Data was analysed using multiple Mann-Whitney U tests with 5% FDR correction. *FDR-corrected discoveries; #Mann-Whitney U test p-value < 0.05 but not an FDR-corrected discovery. No significant comparisons were found between ‘SEA and SA’ groups. B) PCoA displays raw Euclidean distances of lipid concentrations multiplied by PCoA eigenvalues in adults only. Dots represent individual data points. One-way MANOVA was then performed, with the Wilks’ lambda p-value indicated as per the graph. Lines represent clustering at 95% kernel density estimation using Scott’s method. n = 60 for East Asian, n = 24 for South Asian and n = 16 for Southeast Asian for both figures.

**Supplemental Figure 11.**
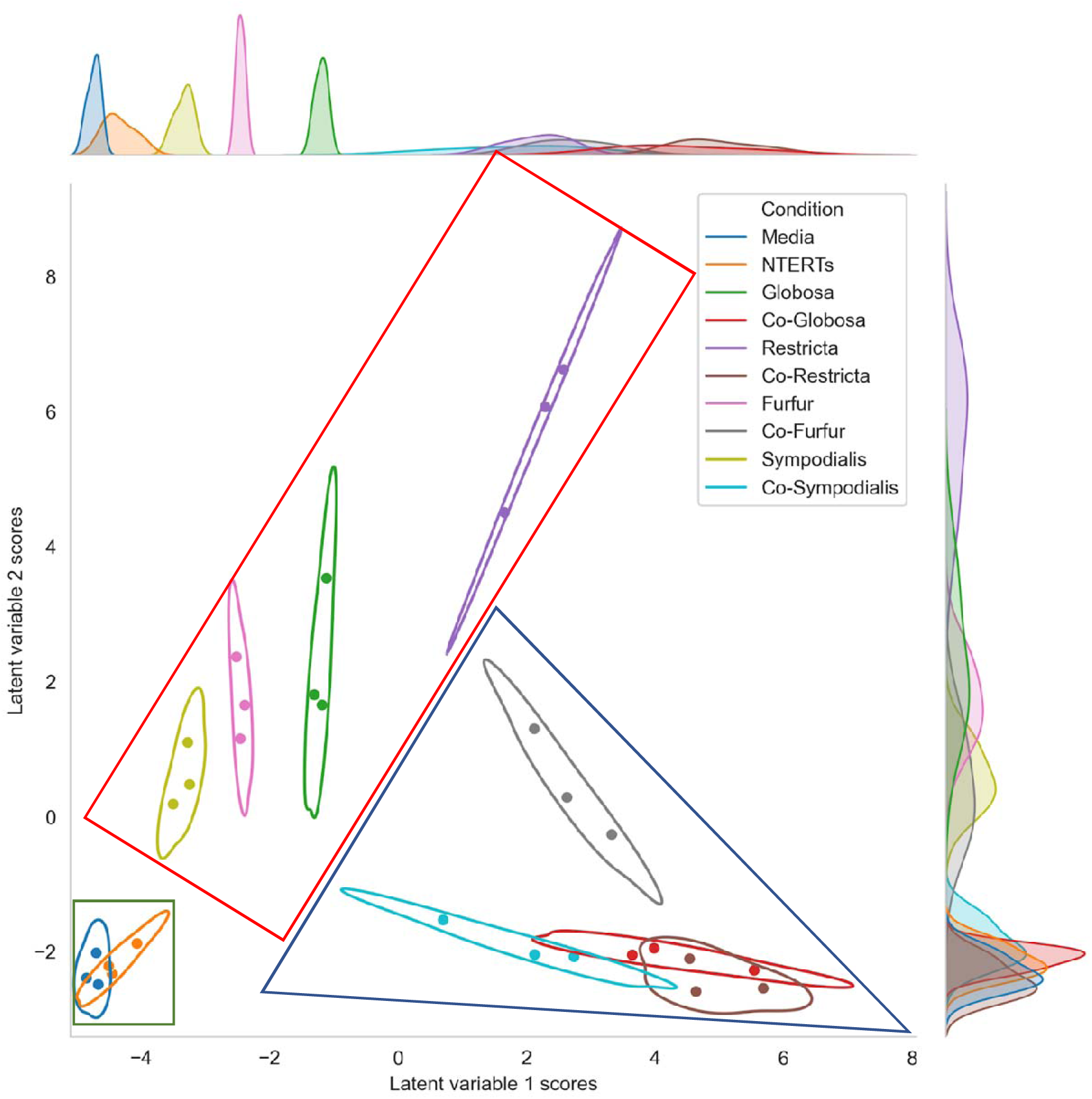
Lipid mediators are different in the various co-cultures. PLS-DA dimensionality reduction was performed with 2 components on every lipid mediator. Dots represent individual data points’ scores in PLS-DA. Lines represent clustering at 95% kernel density estimation using Scott’s method. *Malassezia* grouping was manually highlighted with a red box, co-cultures with a blue triangle, media control and N/TERT-1 mono-culture with a green box.

**Supplemental Figure 12.**
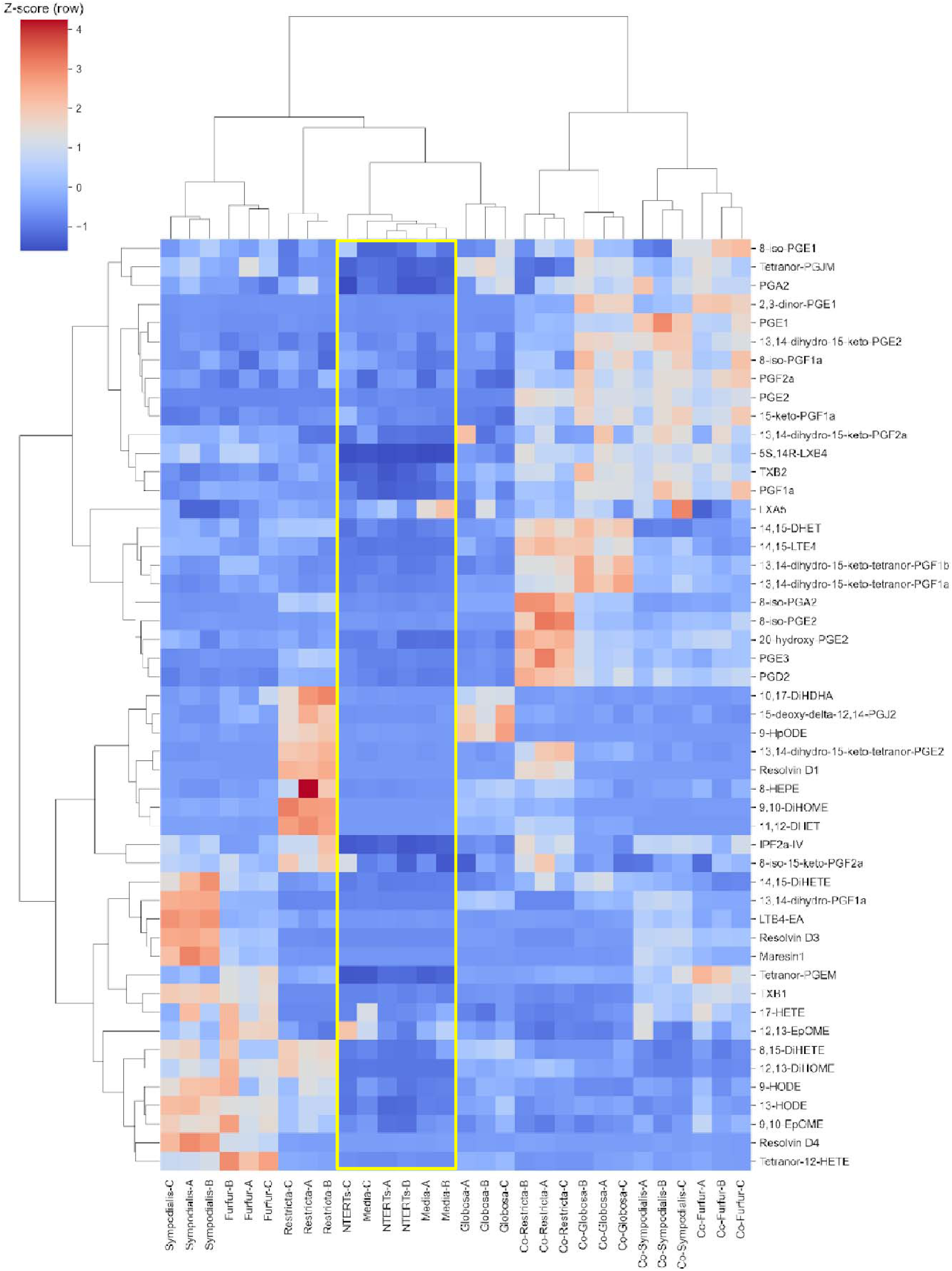
Lipid mediators cluster distinctively in different *in vitro* cultures. To cluster *in vitro* keratinocyte lipid mediators with cytokines, Partial Least Squares - Discriminant Analysis (PLS-DA) with a maximum of 9,999 iterations and a tolerance cut-off of 1e-256 was used in training the model. Z-score, or standard deviation from the mean was then computed per lipid mediator (row-wise) and each subject clustered using Bray-Curtis distance with Farthest Point Algorithm (FPA) hierarchical clustering. Red and blue indicates higher and lower lipid concentration relative to the mean, respectively. Media and N/TERT-1 controls are boxed in yellow for improved visualization.

**Supplemental Figure 13.**
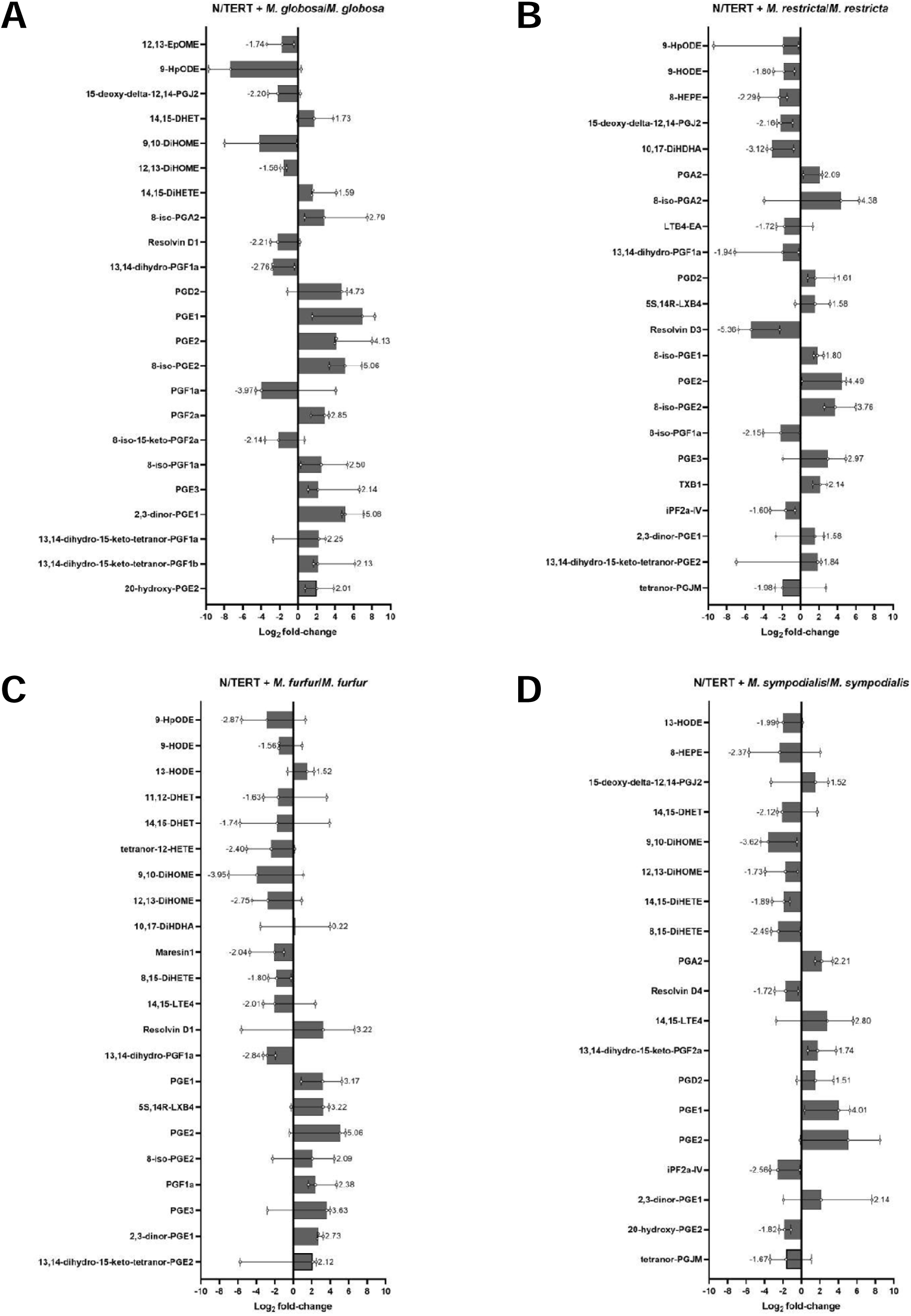
Differences in lipid mediator concentration between N/TERT-1 and *Malassezia* co-cultures versus *Malassezia* mono-cultures. A) N/TERT-1 and *M. globosa* versus *M. globosa* only, B) N/TERT-1 and *M. restricta* versus *M. restricta* only, C) N/TERT-1 and *M. furfur* versus *M. furfur* only, and D) N/TERT-1 and *M. sympodialis* versus *M. sympodialis* only. Only lipid mediators with Log_2_ fold-change ≥ 1.5 between the co-cultures and mono-species cultures are plotted. Bar graphs represents the median of three biological replicates (n = 3) and error bars represent 95% CI.

**Supplemental Figure 14.**
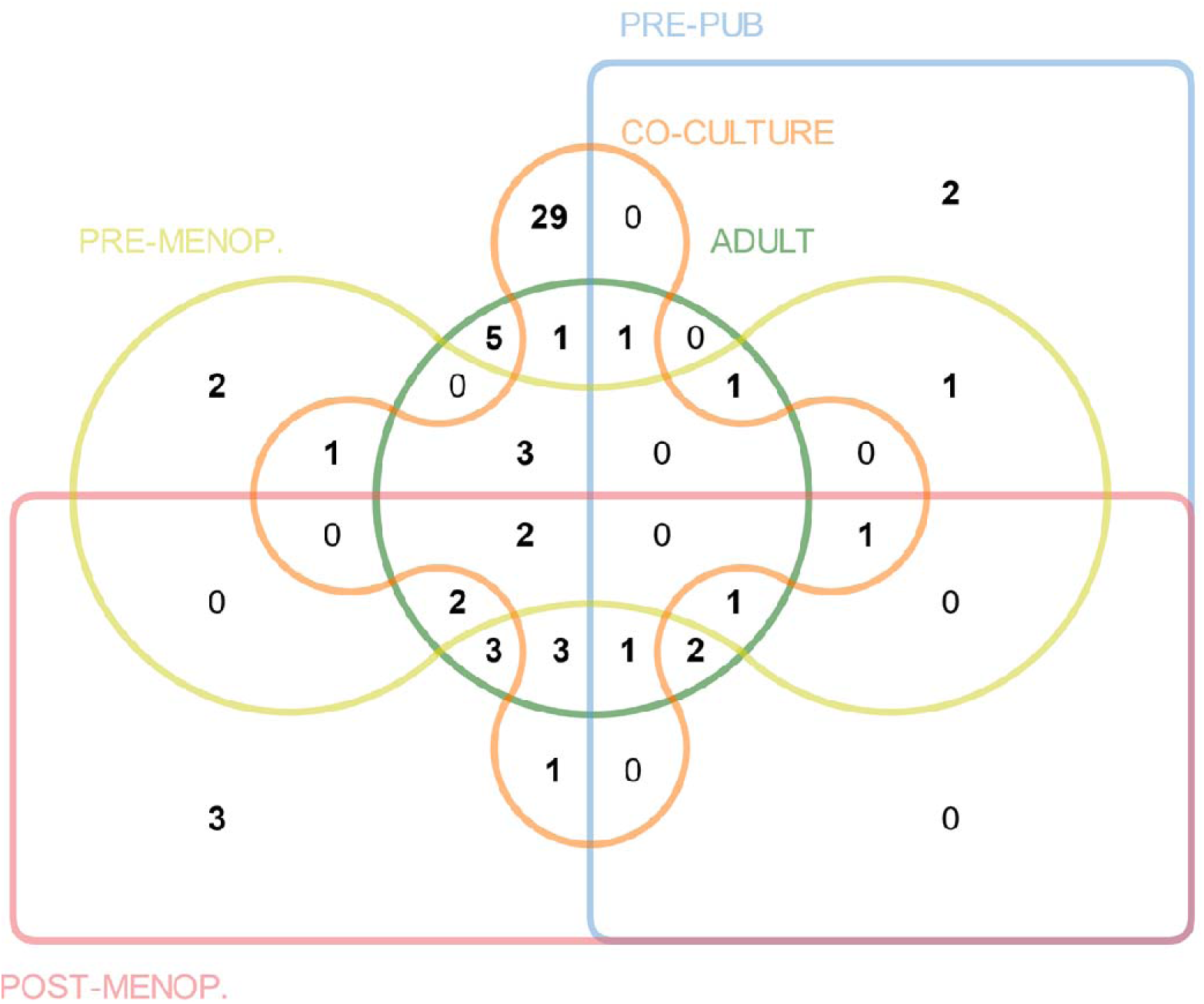
Shared characteristics of skin lipid mediators between subjects from different life stages and *in vitro* co-cultures. Venn diagram shows numbers of overlapping lipid mediator species that are correlated with specific microbiome taxonomies on skin of pre-pubescent, adult, pre- and post-menopausal volunteers and lipid mediator concentrations that are affected by *Malassezia*-keratinocyte co-culture conditions.

